# Integrating Biological and Machine Learning Models for Rainbow Trout Growth: Balancing Accuracy and Interpretability

**DOI:** 10.1101/2025.11.04.686633

**Authors:** Pin Lyu, Lawrence Fulton

## Abstract

Invasive species management demands predictive models that balance accuracy with ecological interpretability. Traditional approaches often fail to capture complex environmental interactions. We evaluated hybrid frameworks integrating biological and machine learning models for rainbow trout (*Oncorhynchus mykiss*) growth in the Lower Colorado River. Using ten years of tag-recapture data and environmental covariates, we assessed traditional and Bayesian von Bertalanffy (VBGM) and Gompertz models, Random Forests, XGBoost, LightGBM, Support Vector Regression, Neural Networks, and ensemble approaches through comprehensive probabilistic comparisons. Our results reveal substantial improvements from incorporating environmental context and advanced modeling. Top methods achieved 70 to 80 percent error reductions compared to baseline models, equivalent to 45 to 70 mm improvements or 20 to 32 percent of mean fish length. A stacked ensemble of XGBoost and the VBGM achieved optimal performance (RMSE = 15.96 mm, *R*^2^ = 0.966), demonstrating complete stochastic dominance across the posterior. Gradient boosting models formed a strong second tier, with LightGBM (9 dominances) and XGBoost (8 dominances) leading this group. Bayesian Model Averaging achieved similar accuracy while explicitly quantifying uncertainty. Even traditional mechanistic models improved markedly, up to 80 percent, when enhanced with covariates and Bayesian estimation. Feature importance analysis identified (on average) initial length, time at large, and weight at release as key predictors. The stacked ensemble dominated baseline models in over 99 percent of posterior samples, confirming its robustness. These findings establish hybrid ensemble frameworks as powerful tools for ecological forecasting, uniting predictive performance with mechanistic insight critical to conservation decision-making. The methodology provides a generalizable template for ecological systems where both accuracy and interpretability are essential.

**Graphical Abstract:** 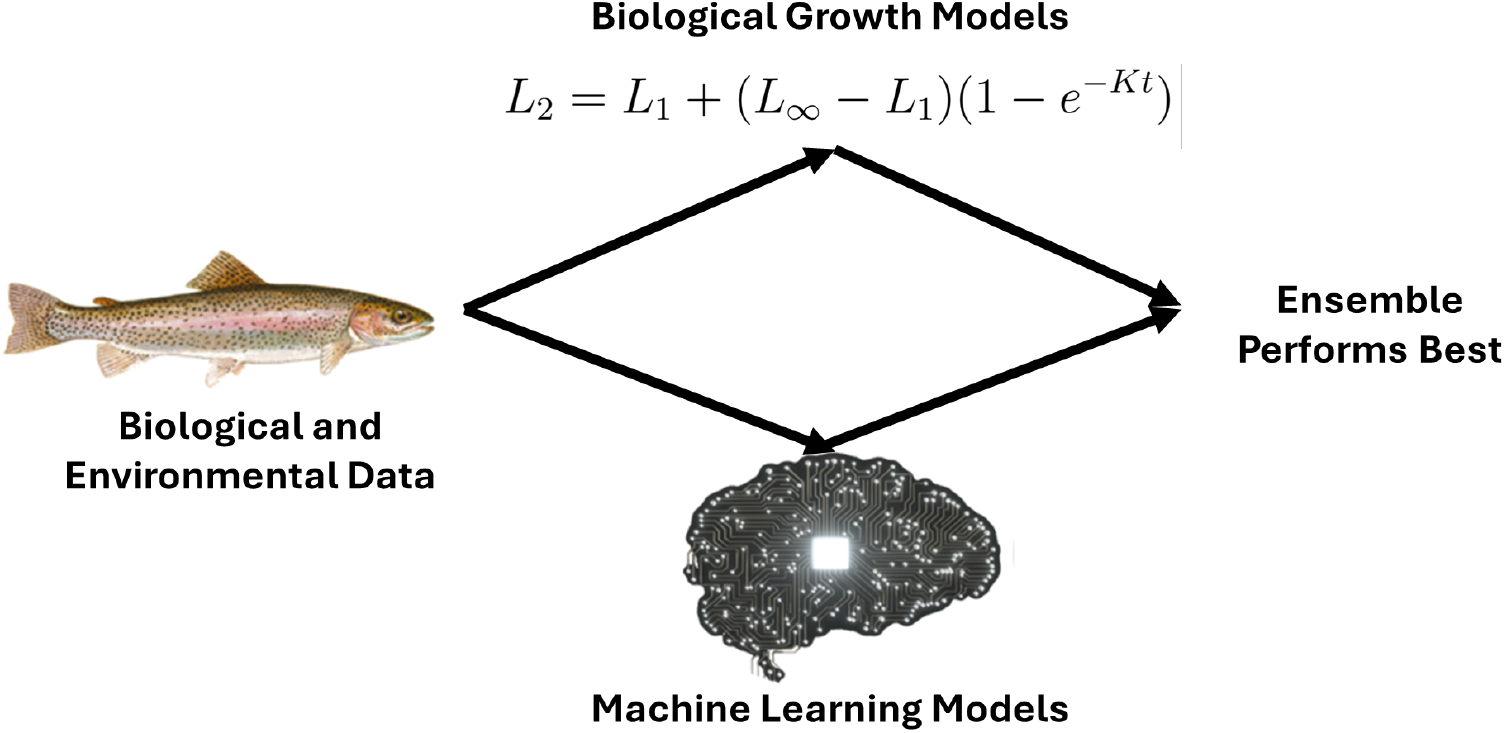

## 1. Introduction

The management of invasive species represents one of the most urgent conservation challenges of the twenty-first century given their profound and escalating ecological and economic impacts. As emphasized by the World Conservation Union, invasive alien species rank among the leading drivers of biodiversity loss and species extinctions worldwide. Their influence extends beyond ecological disruption. Diagne et al. (2021) estimated that biological invasions imposed a minimum financial burden of $1.288 trillion USD on the United States economy between 1970 and 2017 alone. The magnitude of these losses underscores the pressing need for innovative, data-driven management strategies capable of addressing the dual threats to ecosystems and economies.

Rainbow trout (*Oncorhynchus mykiss*), a species native to the Pacific Northwest of the United States, exemplifies the complex trade-offs inherent in invasive species management. Introduced intentionally into the lower Colorado River basin from 1964 to 1998 to support recreational sport fishing (Korman, 2009), rainbow trout have flourished under the cold-water conditions created by Glen Canyon Dam releases. Their rapid expansion, compounded by a lack of natural predators, has led to significant competitive displacement of native endangered fishes such as the humpback chub (*Gila cypha*) and razorback sucker (*Xyrauchen texanus*). Empirical evidence demonstrates that reductions in rainbow trout density are correlated with improved growth rates among native fish, highlighting the disruptive ecological role that this species now plays (Yard et al., 2016).

Paradoxically, rainbow trout also represent a major economic asset. In 2020 alone, the Arizona Game and Fish Department (AZGFD) sold 273,902 fishing licenses, generating nearly $14 million to fund conservation programs. The broader recreational fishing industry contributed over $1.4 billion to the state’s economy, with anglers dedicating more than six million fishing days statewide. Managing rainbow trout populations thus requires navigating a delicate balance between protecting the ecological integrity of the Colorado River ecosystem and sustaining significant economic benefits tied to recreational fisheries (Congressional Sportsmen’s Foundation, 2021).

Growth is central to invasive species management because it determines competitive dynamics, reproduction timing, and the potential for ecological disruption. This study addresses an important need by integrating biologically grounded growth models with modern machine learning (ML) techniques to forecast rainbow trout fork length growth across the Lower Colorado River basin. Unlike prior efforts focused solely on mechanistic or black-box models, our approach fuses biological interpretability with state-of-the-art predictive accuracy, offering both ecological insight and actionable forecasts for targeted intervention

Effective management of this complex ecological and economic challenge requires accurate growth prediction models that can inform targeted interventions. However, existing approaches face inherent trade-offs. Traditional biological models provide interpretable parameters essential for management decisions but may lack flexibility to capture complex environmental interactions, while ML methods excel at modeling nonlinear relationships but offer limited ecological insight. To address this gap, this study integrates the biological interpretability of mechanistic growth models with the predictive flexibility of ML techniques to forecast rainbow trout growth while preserving the ecological understanding essential for adaptive management.

The resulting framework provides resource managers with a transferable set of tools to anticipate population dynamics under varying environmental and policy conditions. By linking individual growth trajectories to broader ecological outcomes, this study demonstrates how hybrid modeling approaches can support adaptive management strategies that balance conservation priorities with socioeconomic interests. Beyond rainbow trout management, these methods offer a generalizable template for ecological forecasting where both predictive accuracy and biological interpretability are essential. Ultimately, the findings provide guidance for evidence-based policymaking in systems where ecological resilience and species management are tightly intertwined.

## 2 Literature Review

Biological growth patterns represent complex nonlinear interactions between environmental drivers and intrinsic factors. Effective growth modeling therefore requires systematic identification of important variables across both exogenous (e.g., water temperature, food availability, competition) and endogenous (e.g., metabolic rate, genotype) domains to parameterize system dynamics (Dutta, 1994).

### 2.1. Determinants of Rainbow Trout Growth

Previous research has identified several variables associated with rainbow trout growth. Sumpter (1991) demonstrated that interactions among photoperiod, water temperature, and genotype affect growth rates through hormonal and metabolic pathways. In addition, trout population density significantly modulates growth outcomes. Korman et al. (2015) found that trout reared at 80 kg/m^3^ exhibited 22% slower growth than those at 40 kg/m^3^, even under ad libitum feeding conditions, attributing the reduction to elevated cortisol levels and intensified competition for food.

Water flow has also been shown to exert complex influences on rainbow trout growth and survival. Korman et al. (2011) highlighted the contrasting effects of High Flow Experiments (HFEs) in the Colorado River. The March 2008 HFE quadrupled juvenile trout survival by enhancing habitat and food availability, while the November 2004 HFE resulted in a threefold decline in age-0 trout, likely due to displacement and mortality. Similarly, Harvey et al. (2006) experimentally reduced streamflow by 75 to 80% in controlled stream sections and observed that rainbow trout in natural flow conditions grew 8.5 times more than those in reduced-flow sections, underscoring the critical role of hydrological conditions in shaping growth outcomes. Nutrient availability, particularly phosphorus (P), further influences growth trajectories. Both dietary phosphorus deficiency and excess have detrimental effects on rainbow trout fry development (Fontagné et al., 2009). Rodehutscord (1996) established that a dietary ratio of 0.25 g available phosphorus per MJ digestible energy is required for optimal trout growth, emphasizing the necessity of precise nutrient management in aquaculture and wild populations.

### 2.2. Foundations of Biological Growth Modeling

Theoretical models of biological growth have evolved to accommodate complex field data structures. Fabens (1965) adapted the von Bertalanffy (1938) growth model (VBGM) to tagging-recapture studies, reformulating it to account for time-at-large between measurements and enabling direct parameter estimation from mark-recapture data. Parallel developments validated the Gompertz growth model, originally proposed for human mortality by Gompertz (1825), as a robust framework for modeling asymptotic growth patterns across a wide range of animal taxa.

Both the VBGM and Gompertz models remain foundational tools in ecological and fisheries science for modeling organismal growth due to their biologically meaningful parameters. These models explicitly incorporate concepts such as asymptotic maximum size and intrinsic growth rate, offering a transparent connection between mathematical formulation and biological reality. This interpretability is particularly valuable in applied management contexts where parameters can be linked to physiological or environmental constraints, such as habitat quality, food availability, or temperature-dependent growth ceilings. Moreover, their compact parameterizations facilitate communication with stakeholders and integration into larger population dynamics models.

However, despite these advantages, both models rely on relatively rigid, predefined functional forms that impose assumptions about the shape and trajectory of growth. For example, VBGM presumes decelerating growth toward a fixed asymptote, while the Gompertz model assumes a sigmoid curve with an inflection point determined by a fixed proportion of the asymptotic size. These forms may not fully capture the variability and plasticity observed in real-world growth trajectories, particularly in dynamic environments where growth can be episodic, nonlinear, or influenced by unobserved factors such as competition, predation risk, or management interventions.

Such rigidity can lead to model misspecification or underfitting, especially when applied to heterogeneous populations like invasive species that exhibit substantial phenotypic flexibility. In the case of rainbow trout in the Lower Colorado River, for instance, individuals may experience rapid early growth due to favorable flow regimes or stocking histories, followed by abrupt slowdowns due to density-dependent effects or seasonal changes. Capturing this heterogeneity requires more flexible modeling approaches or hybrid frameworks that preserve the interpretability of classical models while adapting to empirical complexity. This recognition motivates the integration of mechanistic and data-driven approaches as a way to retain biological relevance while improving predictive performance.

### 2.3. Machine Learning Applications in Biological Growth Prediction

In recent years, ML has emerged as an important and increasingly transformative tool in fisheries science and aquatic ecology. ML techniques can flexibly capture complex, nonlinear relationships between growth outcomes and multiple predictor variables without requiring predefined functional relationships, making them particularly attractive for biological growth modeling.

Ho and Goethals (2022) demonstrated the value of ML for water quality prediction in freshwater ecosystems, while Flores et al. (2024) and Isguzar et al. (2024) successfully applied deep learning methods to predict reproductive status and estimate fish age, respectively. In a related study, Li et al. (2021) applied ML algorithms to predict fish growth patterns, validating their superior predictive accuracy over traditional parametric models.

Beyond fisheries, broader ecological studies have illustrated ML’s effectiveness in biological growth modeling. For example, Pineda-Metz et al. (2023) employed artificial neural networks to model oyster growth under variable environmental conditions, achieving substantially higher accuracy than standard growth models. Similarly, Bhai et al. (2024) used gradient-boosting methods to predict plant growth parameters under diverse environmental regimes, illustrating ML’s cross-domain applicability to biological growth processes.

Despite these advances, relatively few studies have systematically applied ML to predict the growth of invasive species in dynamic riverine ecosystems, particularly for species like rainbow trout (*Oncorhynchus mykiss*), which are both economically valued and ecologically disruptive. Even more notably, the explicit ensembling of mechanistic biological models and ML techniques remains largely absent from the published literature in the ecology discipline. This study addresses that gap. By combining the biological interpretability of traditional models with the flexible accuracy of ML, the resulting ensemble framework not only improves predictive performance but also contributes a novel methodological advance. This approach provides a powerful and adaptable tool for guiding invasive species management, supporting native species recovery, and informing long-term conservation policy in complex and changing freshwater systems.

## 3. Methods

### 3.1. Data, Preprocessing, and Variable Definitions

#### 3.1.1. Data and Limitations

The data used in this study are from the United States Geological Survey (USGS) datasets, *Rainbow Trout Growth Data* and *Growth Covariate Data* from Glen Canyon, Colorado River, Arizona (2012–2021) (Korman and Yard, 2017). These freely-available datasets include ten years of rainbow trout growth measurements from release and recapture surveys in the lower Colorado River basin, along with seven environmental covariates known to influence growth rates. We have also posted all data and code online for replication.

The dataset originates from a tag-recapture program in which rainbow trout were physically tagged upon initial capture. However, a limitation is the absence of consistent individual identifiers across capture events. While tagging was conducted, the dataset does not include reliable codes linking multiple observations of the same fish. As a result, some individuals may appear in the dataset more than once without a definitive way to track their recapture history. Given this constraint, we treated each capture event as an independent observation with its associated covariates (e.g., age, environmental factors, location). This decision avoids imposing assumptions about individual identity but introduces a potential for pseudo-replication if some fish are included multiple times. Because we cannot distinguish repeat captures from new individuals, within-individual temporal dependence was not explicitly modeled. Future studies could improve upon this by ensuring traceable tag identifiers and applying longitudinal or hierarchical models to fully leverage the repeated-measures structure when available.

To accommodate the temporal structure of the data in a biologically meaningful way, we modeled the time between release and recapture events using each fish’s specific *time-at-large*, the duration (in days) between capture events. Rather than using static calendar timepoints, we leveraged interval-based alignment of biometric and environmental data. Environmental variables were recorded as monthly means between survey trips, while fish data included both release and recapture dates. We applied two integration protocols: (1) For fish recaptured within a single monthly interval, the corresponding monthly mean values were assigned to the observation. (2) For fish spanning multiple months, we calculated time-weighted averages of environmental variables over the entire time-at-large period. This approach ensured that covariates reflected the fish’s actual environmental exposure, preserving both ecological fidelity and temporal accuracy in model development.

Although the lack of individual IDs precluded the use of formal mixed-effects or autocorrelation models, our design mitigates these concerns by encoding relevant temporal dynamics through biologically informed durations and matched covariates. The final dataset included 9,798 observations of catch and release with complete environmental integration. No observations were excluded. Many of the algorithms we employed (Random Forest, XGBoost, Neural Networks) are robust to mild violations of independence assumptions and can accommodate the potential pseudo-replication without strong parametric assumptions. Bayesian methods provide a principled framework for explicitly modeling interdependence through joint probability structures, such as hierarchical models, spatial processes, and temporal autocorrelation structures.

#### 3.1.2. Variables (Features)

Each variable in the merged dataset was selected based on its biological relevance to rainbow trout growth and habitat dynamics. The response variable, *L*_2_ or fork length at recapture, is a direct indicator of somatic growth, a critical metric for understanding individual fitness and population health. Time at large captures the duration over which growth can occur and may interact with seasonal and environmental conditions.

The independent variables used in the growth model fall into five biologically meaningful categories: initial condition, spatial, seasonal, temporal, and environmental. Each category is explicated in order.

Initial condition variables include fork length at release (*L*_1_) and weight at release, These variables are crucial for modeling size-dependent growth trajectories. Smaller individuals often exhibit higher relative growth rates due to elevated metabolic demands and compensatory growth dynamics. These initial traits anchor the baseline from which subsequent growth is measured.

Spatial variation is represented by the river mile at release, which captures fixed geographic differences in habitat quality along the river corridor. These differences can influence early growth potential due to longitudinal gradients in water temperature, prey density, substrate, shading, and flow regimes. River position is especially relevant in riverine systems where upstream and downstream reaches may differ markedly in ecological conditions.

Seasonal effects are captured by both the release month and recovery month (both converted to sets of indicator variables), which represent seasonal cycles in water temperature, photoperiod, and prey availability, particularly aquatic insects. Seasonal timing also relates to high-flow events, which can alter habitat structure and displace prey, thereby influencing energy intake and expenditure.

Temporal exposure is modeled using time at large, which quantifies the duration available for growth between release and recapture. In addition, release year and recovery year (again, sets of indicator variables) account for interannual variability in unmeasured but biologically important conditions such as drought, flood pulses, or shifts in food web dynamics. These temporal covariates help capture broad-scale environmental fluctuations that may not be reflected in short-term or localized metrics.

Environmental variables include average river discharge over the time-at-large interval, water temperature, solar insolation, reactive phosphorus concentration, and rainbow trout biomass. Discharge influences habitat availability, drift-feeding opportunities, and the energetic cost of station-holding. Water temperature governs metabolic rate and food conversion efficiency, constraining growth within species-specific thermal limits. Solar insolation serves as a proxy for in-stream primary production, affecting food web productivity. Reactive phosphorus concentration is a limiting nutrient for algal growth, thereby modulating base-level productivity. Biomass provides an ecologically valid index of density-dependent pressures, such as competition for resources. Rather than relying on individual fish metrics, biomass estimates were derived from a Jolly-Seber mark-recapture model applied to size-class abundance data across the central 3 km of the study reach (Korman and Yard, 2017), reflecting population-level conditions that influence growth trajectories.

#### 3.1.3. Training, Validation, and Test Sets

Following data merging and feature engineering, we randomly split the dataset into training and testing subsets using a 70/30 ratio, with a fixed pseudo-random number seed to ensure reproducibility. Seventy percent of the data was allocated for training, while the remaining 30% served as a held-out test set for evaluating model generalization. To prevent information leakage, no test data was used during model development. Within the training set, we conducted hyperparameter tuning via 5-fold cross-validation. Specifically, we employed randomized search over a targeted hyperparameter space for each ML model to identify optimal configurations while managing computational cost. This strategy balances tuning efficiency with robustness and helps reduce the risk of overfitting. Because model evaluation was performed exclusively on an untouched 30% test set, completely isolated from training and hyperparameter tuning, the resulting performance metrics provide an unbiased estimate of true generalization ability.

#### 3.1.4. Data Scaling

To prepare the data for model training, two scaling strategies were considered to normalize the predictor variables. In the first approach, all continuous predictors were scaled to a [0, 1] range using a Min-Max transformation. This full-scaling strategy was used for ML models, where consistent feature ranges are important for convergence and optimization. In the second approach, two biologically meaningful predictors, *L*_1_ (initial length) and Time at Large, were excluded from scaling to preserve their native units and biological interpretability. This partial-scaling strategy was applied to models grounded in biological reasoning. All remaining continuous features in this approach were scaled identically to the full-scaling method. In both cases, scaling parameters were computed using only the training data and then applied to the test data, ensuring that no information from the test set leaked into the model development process. Importantly, the response variable (*L*_2_) was never scaled to support interpretability. Both scaled datasets were used in parallel to assess whether preserving the original scale of biologically relevant variables improved interpretability or predictive performance.

#### 3.1.5. Software

All data preprocessing, model implementation, and statistical analyses were conducted using **Python 3.13**, the latest stable release of the Python programming language (Python Software Foundation, 2025). Python offers a robust open-source coding ecosystem ideal for ML, data wrangling, and statistical analysis, with libraries such as NumPy (Harris et al., 2020), pandas (McKinney et al., 2010), scikit-learn (Pedregosa et al., 2011), XG-Boost(Chen and Guestrin, 2016a), LightGBM (Ke et al., 2017), and Pytorch (Paszke et al., 2019), enabling efficient and reproducible modeling workflows. Python’s widespread adoption in the scientific community, flexible syntax, and extensive package repository made it an ideal platform for this research.

## 4. Models

To model rainbow trout growth from the available data, we employed both biologically grounded growth models, contemporary ML approaches, and an ensemble approach. Each modeling paradigm offers distinct and complementary advantages. Biological models, such as the VBGM and its derivatives, yield interpretable parameters, e.g., intrinsic growth rate and asymptotic size, that are directly linked to physiological processes and life-history traits (von Bertalanffy, 1938; Fabens, 1965). These models are essential for understanding mechanistic dynamics and aligning findings with ecological theory.

In contrast, ML models are data-adaptive and excel at uncovering complex, nonlinear relationships that often arise in ecological systems but may be difficult to specify a priori using closed-form equations (Ho and Goethals, 2022; Li et al., 2021). Their flexibility allows for the inclusion of a broad suite of predictors (particularly environmental covariates) without requiring strong assumptions about functional form. Many ML algorithms also provide variable importance metrics, offering insight into the relative influence of each predictor on growth outcomes. This feature is especially valuable in ecological contexts, where interactions among factors such as temperature, discharge, and biomass can be multifaceted and hierarchical.

That said, parametric models such as the VBGM and the Gompertz function can be extended to include covariates. When specified appropriately, these models allow researchers to embed biological knowledge directly into the functional form, offering mechanistic interpretability that complements the more flexibleâĂŤbut often less transparentâĂŤML approaches. We adopted this approach in our analysis, incorporating covariates within these growth models and estimating parameters using a Bayesian framework.

Ensembles integrate predictions from multiple models, either within a single class or across different modeling paradigms, to improve overall performance. For instance, a random forest internally aggregates the outputs of numerous decision trees, each trained on different data subsets and feature selections. In contrast, the framework proposed here represents a heterogeneous ensemble, combining biologically grounded models such as the Bayesian VBGM with data-driven ML algorithms like XGBoost. This cross-model integration allows the ensemble to leverage the complementary strengths of each approach, biological interpretability and flexible predictive power on the other. Such heterogeneity often enhances predictive accuracy by increasing model diversity, which is a critical factor in ensemble success. As Brown et al. (2005) demonstrated, there exists a trade-off between individual model accuracy and ensemble diversity, and optimal performance is often achieved when models make uncorrelated errors while maintaining reasonable base accuracy. By strategically combining mechanistic and ML models, this study not only improves prediction of somatic growth in invasive rainbow trout but also contributes a novel example of the accuracy-diversity principle applied in an ecological forecasting context.

Our selection of modeling approaches reflects these complementary priorities. Our initial benchmarks were the deterministic VBGM and Gompertz without covariates. We then included Bayesian-estimated nonlinear models such as VBGM and the Gompertz function to retain ecological interpretability and biological realism. These were contrasted with ML algorithms chosen to represent a broad set of methodological paradigms: Support Vector Regression (SVR) for kernel-based regularization, artificial neural networks (ANNs) for high-capacity function approximation, and tree-based ensembles (Random Forest, XGBoost, LightGBM) for robust performance, embedded feature selection, and interpretability. We also incorporated a Bayesian linear model to examine how parameter uncertainty and prior regularization influence predictions in comparison to both biological and ML alternatives. This multi-model framework allowed us to evaluate a wide range of modeling assumptions, from mechanistic to data-driven, under consistent data conditions.

### 4.1. Biological Growth Models

This study implemented two biologically motivated growth models: the Fabens version of the VBGM (Fabens, 1965), and the Gompertz growth model (Gompertz, 1825). Both were estimated using Bayesian methods.

The Fabens VBGM formulation follows.

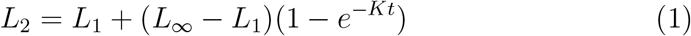

where:

- *L*_2_: fork length at recapture,
- *L*_1_: fork length at initial marking (release),
- *L*_∞_: asymptotic maximum fork length,
- *K*: growth coefficient,
- *t*: time (in years) between marking and recapture.

This formulation captures somatic growth as the combined effect of two forces: the approach toward a genetically or environmentally constrained maximum size (*L*_∞_) and the declining influence of the fish’s initial length over time. This structure is especially suitable for tag-recapture data, where age is unknown but the time interval between observations is well-defined.

Rewriting the model algebraically emphasizes its dual components:

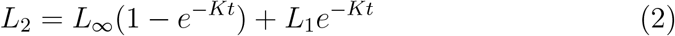

This version illustrates how the observed length at recapture reflects a weighted combination of asymptotic potential and initial condition, with weights governed by the exponential decay term *e*^−*Kt*^.

#### 4.1.1. Bayesian Fabens Model Formulation

To estimate the parameters of the Fabens VBGM using Bayesian inference, we specified the following hierarchical model:

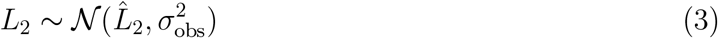

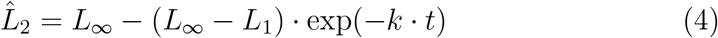

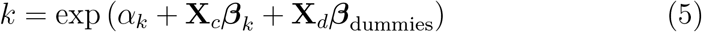

where:

- *L*_1_: fork length at initial marking,
- *L*_2_: fork length at recapture,
- *t*: time at large (in years),
- *L*_∞_: asymptotic maximum fork length,
- *k*: growth rate coefficient (positive, covariate-dependent),
- **X**_*c*_: matrix of scaled continuous covariates,
- **X**_*d*_: matrix of dummy variables for seasonal effects,
- ***β***_*k*_ : vector of regression weights for continuous covariates,
- ***β***_dummies_: vector of regression weights for dummy predictors.

The prior distributions were specified as follows:

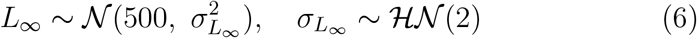

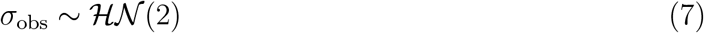

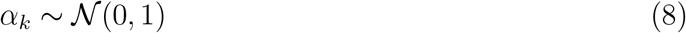

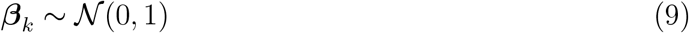

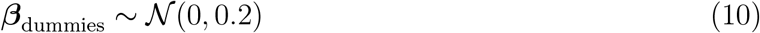

##### Priors

Priors were selected to reflect weakly informative beliefs consistent with known biological constraints and to regularize the estimation of growth parameters in the presence of covariates. The asymptotic length *L*_∞_ was assigned a normal prior centered at 500 mm, with a data-informed standard deviation drawn from a half-normal distribution ℋ𝒩 (2). This reflects prior knowledge that asymptotic sizes for rainbow trout in similar systems typically fall between 400–600 mm, while still allowing flexibility. The observation error *σ*_obs_ and the hyperparameter 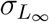 were each given half-normal priors with scale 2, reflecting reasonable uncertainty around measurement noise and biological variability. The intercept and continuous covariate effects on the growth coefficient *k* were modeled with standard normal priors 𝒩 (0, 1), supporting broad exploration of potential influences while enforcing shrinkage toward zero in the absence of signal. Seasonal dummy effects were assigned tighter priors 𝒩(0, 0.2) to prevent overfitting from sparse or collinear binary predictors and to reflect their role as adjustment factors rather than primary drivers. All priors were selected to ensure identifiability and facilitate convergence during sampling, without imposing strong constraints on parameter estimates. The model was implemented in *NumPyro* (Phan et al., 2019), and *posterior samples were obtained via Markov Chain Monte Carlo (MCMC) using the No-U-Turn Sampler (NUTS)(Hoffman et al., 2014)*.

##### Formulation

The joint posterior distribution of the model parameters

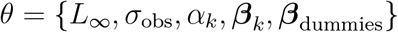

given the observed data

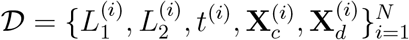

is proportional to the product of the likelihood and prior distributions:

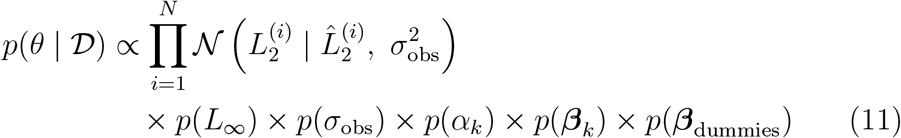

where the predicted length at recapture is defined as:

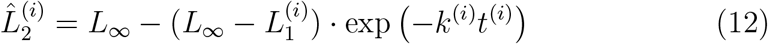

and the individual-specific growth coefficient *k*^(*i*)^ is modeled as:

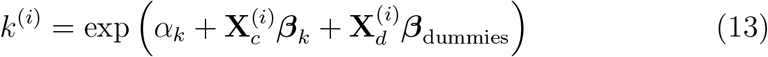

#### 4.1.2. Bayesian Gompertz Model Formulation

The second biological model implemented was the Gompertz (1825) growth equation, a sigmoid-shaped function that describes asymptotic growth using a double-exponential form. Originally developed to model human mortality, the Gompertz function has since been widely applied in biological and ecological contexts to capture growth processes that begin rapidly but decelerate over time as they approach an upper physiological limit. The model assumes that the relative growth rate decreases exponentially with time, making it particularly well-suited for species exhibiting rapid early development followed by gradual slowing as size approaches a maximum threshold. The Gompertz equation models the predicted fork length *L*_2_ at time *t* as:

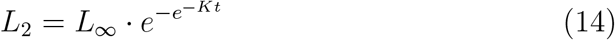

To estimate the parameters of the Gompertz growth model using Bayesian inference, we specified the following hierarchical model:

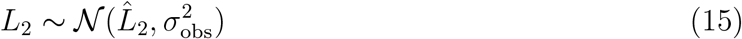

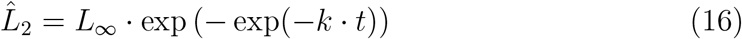

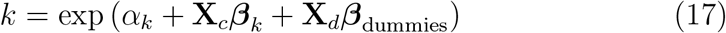

where:

- *L*_2_: fork length at recapture,
- *t*: time at large (in years),
- *L*_∞_: asymptotic maximum fork length,
- *k*: growth rate coefficient (positive, covariate-dependent),
- **X**_*c*_: matrix of scaled continuous covariates,
- **X**_*d*_: matrix of dummy variables for seasonal effects,
- ***β***_*k*_ : vector of regression weights for continuous covariates,
- ***β***_dummies_: vector of regression weights for dummy predictors.

The prior distributions were specified as:

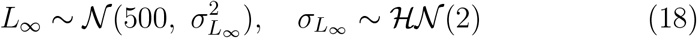

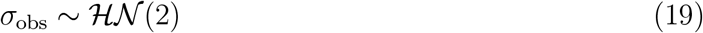

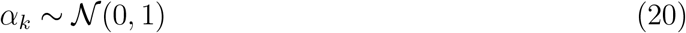

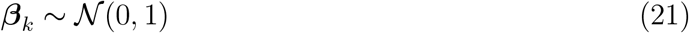

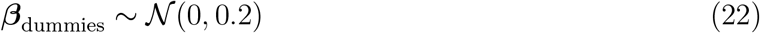

##### Priors

Priors for the Gompertz model were specified identically to those in the Fabens VBGM to ensure consistency and comparability across formulations. All priors reflect weakly informative biological expectations while promoting identifiability. The asymptotic fork length *L*_∞_ was assigned a normal prior centered at 500 mm, with a scale governed by a half-normal distribution ℋ 𝒩 (2), allowing flexibility while anchoring estimates within empirically observed trout size ranges. Both *σ*_obs_ and 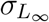 were given half-normal priors to constrain positive variances without imposing rigid assumptions. The intercept *α*_*k*_ and continuous covariate coefficients ***β***_*k*_ were assigned 𝒩 (0, 1) priors to support moderate deviations while avoiding overdispersion. Dummy variable coefficients ***β***_dummies_ were regularized with narrower 𝒩 (0, 0.2) priors to mitigate overfitting from sparse or collinear seasonal predictors. By applying the same prior structure, we ensure that posterior differences between models are attributable to differences in functional form, rather than prior-induced biases. The model was also implemented in *NumPyro* (Phan et al., 2019), and posterior sampling was conducted using NUTS (Hoffman et al., 2014)).

##### Formulation

The joint posterior distribution of the model parameters

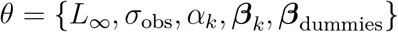

given the observed data

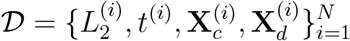

is proportional to the product of the likelihood and prior distributions:

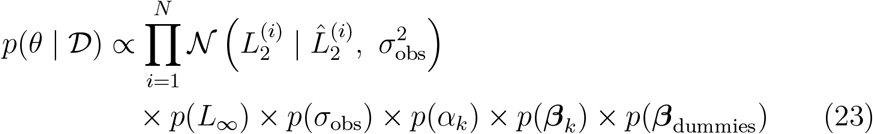

where the predicted length at recapture is defined as:

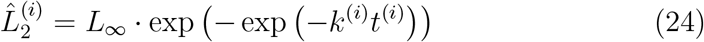

and the individual-specific growth rate is:

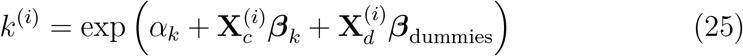

#### 4.1.3. Bayesian Justification

We employed Bayesian inference for both the Fabens VBGM and Gompertz growth models due to its advantages in handling ecological data with complex structure and limited information. Bayesian methods allow for the incorporation of biologically informed prior knowledge–such as plausible bounds for asymptotic size or expected variability in growth rates–which improves parameter regularization and interpretability. These methods do require scaling of the non-indicator independent variables. This is especially critical in hierarchical models with latent variables and covariate-dependent growth rates, where frequentist estimators may yield unstable or biologically implausible results. Additionally, Bayesian models yield full posterior distributions, which enable more comprehensive uncertainty quantification and straightforward propagation of parameter uncertainty through model predictions. The Bayesian framework also supports posterior predictive checks, probabilistic forecasts, and model comparison using information criteria and cross validation tools that are not as readily applicable or interpretable under a frequentist paradigm. Taken together, these features make the Bayesian approach more robust, flexible, and ecologically appropriate for modeling rainbow trout growth under temporally and spatially structured environmental conditions.

### 4.2. Statistical and ML Models

In addition to the biologically grounded growth models, we implemented a diverse suite of statistical and ML algorithms to evaluate predictive performance across modeling paradigms. These included a Bayesian linear model for interpretable, uncertainty-aware regression; a Linear SVR model for capturing linear relationships with regularization (Drucker et al., 1996); a kernel-optimized SVR (with permutative importance), and several nonlinear, tree-based ensemble methods, including Random Forest (RF) (Breiman, 2001), XGBoost (Chen and Guestrin, 2016b), and LightGBM (LGBM) (Ke et al., 2017), each known for their robustness, embedded feature selection, and scalability. We also implemented an Artificial Neural Network (ANN) (McCulloch and Pitts, 1943), representing a high-capacity function approximator capable of modeling complex, nonlinear relationships. Finally, to leverage the complementary strengths of mechanistic and data-driven approaches, we constructed two heterogeneous ensembles that combined the best-performing biological model with the top-performing ML model. This ensemble framework was designed to enhance predictive accuracy while preserving biological interpretability, aligning with recent advances in integrative ecological modeling.

#### 4.2.1. Bayesian Linear Model Formulation

We implemented a Bayesian linear model to serve as a baseline for evaluating growth prediction under additive linear assumptions. The model assumes a normal likelihood with a linear predictor and constant variance:

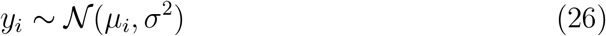

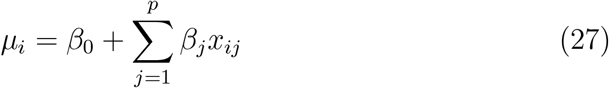

where:

- *y*_*i*_: fork length at recapture (response variable),
- *x*_*ij*_: value of the *j*th predictor for observation *i*,
- *β*_0_: model intercept,
- ***β*** = *{β*_1_, …, *β*_*p*_*}*: regression coefficients,
- *σ*: observation noise (residual standard deviation).

We placed weakly informative priors on all regression parameters to stabilize estimation without imposing strong constraints. The regression coefficients were assigned independent normal priors:

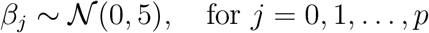

and the standard deviation of residuals was parameterized as:

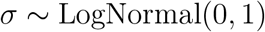

with a deterministic transformation for interpretability:

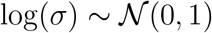

We chose a Bayesian approach for the linear model to maintain consistency across other modeling frameworks and to take advantage of posterior inference, particularly under uncertainty and moderate sample sizes. Bayesian linear regression provides full posterior distributions for all parameters, allowing direct probabilistic interpretation and better uncertainty quantification than frequentist confidence intervals. Additionally, posterior draws for regression coefficients enable direct comparison with effect estimates from the nonlinear growth models. This formulation also improves robustness to multicollinearity and overfitting in high-dimensional predictor sets through regularization induced by priors. Unlike frequentist ordinary least squares (OLS), which provides only point estimates and asymptotic standard errors, the Bayesian version yields complete joint distributions and facilitates posterior predictive checks and model averaging. Moreover, by expressing the observation error in log-space, we achieved more stable sampling and better convergence behavior in MCMC, particularly under heteroscedastic conditions. The model was implemented in *NumPyro*, and posterior samples were drawn using NUTS (Hoffman et al., 2014).

#### 4.2.2. Random Forest (RF)

An RF (Breiman, 2001) is an ensemble learning method that constructs a collection of decorrelated decision trees and aggregates their predictions to produce a more stable and accurate model. For regression tasks, the predicted outcome *ŷ* is computed as the average of *T* individual decision tree predictions:

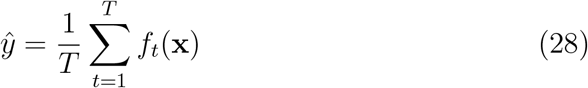

where:

- *T* is the number of trees in the forest,
- *f*_*t*_(**x**) is the prediction of the *t*-th tree.

Each decision tree is trained on a bootstrap sample drawn with replacement from the original dataset, a process known as *bagging* (bootstrap aggregating). Furthermore, at each node split, RF consider only a random subset of predictors, introducing additional decorrelation between trees and reducing overfitting.

By averaging over many uncorrelated trees, RF significantly reduces the variance of individual tree estimators. While single decision trees are prone to high variance and may overfit training data, the ensemble average acts as a variance stabilizer, yielding a more generalizable model. This property is particularly advantageous in ecological datasets, which often contain complex interactions, nonlinearities, and noisy measurements. Each individual tree uses a greedy algorithm to recursively partition the feature space based on impurity measures such as Mean Squared Error (MSE) used in this study.

Importantly, RF offer a built-in mechanism for assessing feature importance. This importance is typically calculated (as it is here) based on the average decrease in impurity (or prediction error) resulting from splits on each feature, aggregated across all trees.

To optimize the performance of the Random Forest model, we employed a randomized hyperparameter search using 5-fold cross-validation. The search focused on key parameters that influence model complexity and generalization, including the number of trees, maximum depth of each tree, the minimum number of samples required to split an internal node, the minimum number of samples required to form a leaf, and the strategy used to select input features at each split. Rather than exhaustively evaluating all possible combinations, we sampled 25 random configurations from the defined hyper-parameter space to balance efficiency and thoroughness. Model performance was assessed using negative mean squared error as the scoring metric, and the configuration yielding the lowest average validation error was selected as the final model.

The Random Forest model was tuned using a randomized search across the following hyperparameter space, with the selected values shown in **bold**:

- Number of trees (`n_estimators`): [100, **200**, 300, 400]
- Maximum tree depth (`max_depth`): [15, **25**, None]
- Minimum samples required to split a node (`min_samples_split`): [**2**, 5, 10]
- Minimum samples required to form a leaf node (`min_samples_leaf`): [**1**, 3, 5]
- Feature sampling method for tree construction (`max_features`): [**sqrt**, log2]

These settings were selected based on performance using randomized search with 5-fold cross-validation, optimizing for generalization and predictive accuracy.

#### 4.2.3. XGBoost and LightGBM (LGBM)

Both XGBoost (Chen and Guestrin, 2016b) and LGBM (Ke et al., 2017) are gradient-boosted decision tree (GBDT) frameworks that build powerful predictive models through the sequential addition of weak learners (typically shallow regression trees). The prediction function is constructed as a sum of *T* individual tree functions:

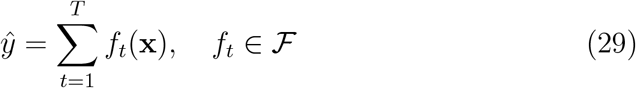

where ℱ is the space of regression trees, and each *f*_*t*_ represents a base learner trained to predict the residuals of the current ensemble.

Unlike bagging-based methods like RF, boosting operates sequentially, where each new tree corrects the prediction error made by the ensemble so far. Both XGBoost and LGBM use gradient descent to minimize an objective function that combines a differentiable loss function ℒ(*ŷ, y*) (typically squared error for regression) with a regularization term Ω(*f*) to penalize model complexity:

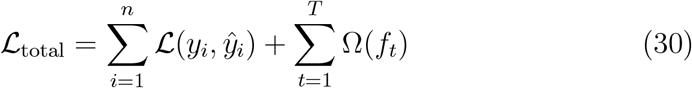

Regularization discourages overfitting and promotes generalization by penalizing trees with excessive depth, number of leaves, or leaf weights.

XGBoost (Chen and Guestrin, 2016a) constructs trees *depth-wise*, growing each level of the tree simultaneously and splitting all nodes at the current depth before proceeding. This strategy produces balanced trees and generally leads to faster convergence with fewer splits.

LGBM (Ke et al., 2017), in contrast, adopts a *leaf-wise* growth strategy: it selects the leaf with the largest reduction in loss and continues splitting from that point. This often results in deeper, more asymmetric trees, allowing for greater flexibility and improved computational efficiency on large datasets. However, it can increase the risk of overfitting on smaller datasets if not properly regularized.

To fine-tune the XGBoost model, we conducted a randomized hyperparameter search using 5-fold cross-validation. The search targeted parameters that directly influence model flexibility, learning dynamics, and regularization. These included the learning rate, maximum tree depth, number of boosting rounds, subsampling ratio for training instances, proportion of features used per tree, and both *L*_1_ and *L*_2_ regularization strengths. Given the use of higher learning rates, the number of boosting rounds was kept moderate to avoid overfitting. We sampled 25 random combinations from the defined hyperparameter space to ensure efficient yet thorough exploration. Model performance was evaluated based on negative mean squared error, and the configuration yielding the lowest average validation error was selected as the final model.

The XGBoost model was tuned using randomized search with 5-fold CV over the following hyperparameter space. The selected values are shown in **bold**:

- Learning rate (`eta`): [**0.05**, 0.1, 0.15, 0.2]
- Maximum tree depth (`max_depth`): [3, **4**, 5]
- Number of boosting rounds (`n_estimators`): [**200**, 400, 600, 800]
- Subsample ratio of training instances (`subsample`): [0.7, **0.8**, 0.9]
- Proportion of features used per tree (`colsample_bytree`): [0.8, 0.9, **1.0**]
- L1 regularization term on weights (`reg_alpha`): [**0**, 0.1]
- L2 regularization term on weights (`reg_lambda`): [**1**, 1.5]

These parameters were selected to balance generalization and computational efficiency, with model performance validated using cross-validation.

For the LightGBM model, we also implemented a randomized hyperparameter search using 5-fold cross-validation. The search was designed to explore parameters central to boosting performance and model regularization. These included the number of boosting rounds, learning rate, maximum tree depth, and the number of leaves per tree, which is particularly influential in LightGBM’s gradient-based framework. Additional parameters such as sub-sampling ratios for rows and columns, along with *L*_1_ and *L*_2_ regularization strengths, were also tuned to improve generalization. A total of 20 random configurations were sampled from the defined hyperparameter space to achieve an efficient and targeted search. Model performance was evaluated using negative mean squared error, and the configuration with the lowest average validation error was selected as the final model.

The LightGBM model was tuned using randomized search across the following hyperparameter space. Selected values are shown in **bold**:

- Number of boosting rounds (`n_estimators`): [100, **150**, 200]
- Learning rate (`learning_rate`): [**0.05**, 0.1, 0.15]
- Maximum tree depth (`max_depth`): [15, **20**, -1]
- Number of leaves per tree (`num_leaves`): [**31**, 50, 100]
- Subsample ratio of training data (`subsample`): [0.7, 0.8, **0.9**]
- Column sampling ratio per tree (`colsample_bytree`): [0.8, **0.9**, 1.0]
- L1 regularization term on weights (`reg_alpha`): [0, **0.1**]
- L2 regularization term on weights (`reg_lambda`): [**0**, 0.1, 1]

#### 4.2.4. Support Vector Regression (SVR)

SVR extends the foundational concepts of Support Vector Machines (SVMs) to regression tasks by constructing an optimal predictive function that balances model complexity with tolerance for small prediction errors (Drucker et al., 1996). SVR estimates a continuous response variable *ŷ* using a linear or nonlinear mapping:

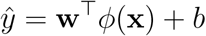

where *ϕ*(·) is a (possibly nonlinear) transformation of the input vector **x, w** is the weight vector, and *b* is the bias term.

Unlike OLS, which minimizes squared error, SVR introduces an *ε*-insensitive loss function that ignores prediction errors within a margin of *ε*. The optimization objective is to find a “flat” function that approximates the data within this margin while minimizing complexity. Formally, the primal optimization problem follows.

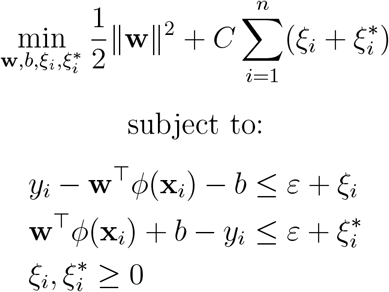

Here, ∥ **w** ∥ ^2^ penalizes model complexity to promote generalization. The slack variables *ξ*_*i*_ and 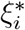 capture deviations beyond the *ε*-tube, and the regularization parameter *C* governs the trade-off between model flatness and tolerance to errors exceeding *ε*.

This convex quadratic programming formulation is typically solved in the dual using Lagrangian multipliers, which naturally identifies a subset of *support vectors*–data points that lie outside the *ε*-tube and directly influence the regression function.

Solving the dual problem enables the use of *kernel functions*, defined as *K*(**x**_*i*_, **x**_*j*_) = *ϕ*(**x**_*i*_)^⊤^*ϕ*(**x**_*j*_), to compute inner products in high-dimensional feature spaces without explicit transformation. This *kernel trick* allows SVR to model nonlinear relationships efficiently. Common choices include the radial basis function (RBF), polynomial, and sigmoid kernels.

To develop an SVR with interpretable coefficients, we initially restricted the hyperparameter search to a linear kernel. This choice facilitates straight-forward interpretation of the feature effects on the predicted outcome and allows comparison with traditional parametric models. While linear regression models also offer interpretable coefficients, we included the linear-kernel SVR model to assess whether a margin-based linear method with regularization could improve generalization performance without sacrificing interpretability. A randomized hyperparameter search with 5-fold cross-validation was employed to tune the penalty parameter, the epsilon-insensitive loss margin, and the kernel coefficient. Model performance was evaluated using negative root mean squared error (RMSE), and the configuration with the lowest average validation error was selected as the final model.

The linear Support Vector Regression (SVR) model was tuned (5-fold CV) using a predefined hyperparameter space. Since a linear kernel was used, the `gamma` parameter was not applicable. The hyperparameter search space and selected values are shown below, with selected values in **bold**:

- Penalty parameter (*C*): [0.1, 1, 10, **100**]
- Epsilon in the loss function (***ϵ***): [0.01, 0.1, **0.5**]

These parameters were selected based on performance using cross-validation, with the linear kernel fixed.

A second Support Vector Regression (SVR) model was formulated to evaluate the optimal kernel type as well as the other parameters. The investigated components mirrored those of the linear SVR, with the addition of kernel selection. Three kernels (linear, polynomial, and radial basis function or RBF) were examined using hyperparameter tuning and 5-fold cross-validation. A nonlinear Support Vector Regression (SVR) model was tuned to evaluate kernel performance, including linear, polynomial, and radial basis function (RBF) kernels. The selected configuration used an **RBF kernel**, with other hyperparameters optimized via randomized search and 5-fold cross-validation. The hyperparameter space and selected values are listed below, with selected values in **bold**:

- Kernel type (`kernel`): [linear, poly, **rbf**]
- Penalty parameter (*C*): [0.1, 1, 10, **100**]
- Epsilon in the loss function (***ϵ***): [**0.01**, 0.1, 0.5]
- Gamma (***γ***): [**scale**, 0.01, 0.1]

This configuration was found to provide the best trade-off between model complexity and predictive performance on the validation folds. To compare the parameter relevance of the SVR model against other ML approaches, **permutative feature importance** was evaluated. This method allowed for a consistent framework to assess the relative contribution of each input variable across models, enabling a robust comparison of feature influence in the presence of nonlinearities and interactions.

#### 4.2.5. Artificial Neural Network (ANN)

Artificial Neural Networks (ANNs) (McCulloch and Pitts, 1943) are layered computational models inspired by the architecture of the human brain. They are capable of approximating highly nonlinear, hierarchical relationships between inputs and outputs through learned feature transformations. In their most basic form, ANNs consist of an input layer, one or more hidden layers, and an output layer, with each layer composed of nodes (neurons) that compute weighted combinations of their inputs. Although artificial neural networks (ANNs) are not always the top-performing choice for structured tabular data, especially when sample sizes are moderate, their inclusion in this study was motivated by both methodological and ecological considerations. From a methodological perspective, ANNs serve as a flexible, universal function approximator capable of capturing complex, nonlinear relationships that may elude simpler parametric models. Given the known nonlinearities in fish growth and environmental interactions, this capacity warranted their inclusion. Ecologically, neural networks allow us to test whether non-tree-based nonlinear models can uncover complementary patterns in rainbow trout growth dynamics, especially in interaction-heavy scenarios where traditional models may oversimplify relationships. Including the ANN also provided a useful baseline for comparing deep learning performance against more interpretable models such as tree ensembles and biologically grounded equations.

The general forward pass for an *L*-layer feedforward neural network is expressed as:

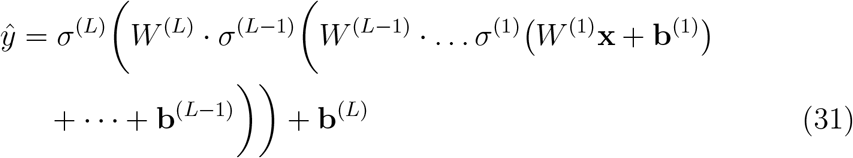

where:

- *W* ^(*l*)^ denotes the weight matrix at layer*l*,
- *l*, **b**^(*l*)^ is the bias vector at layer *l*,
- *σ*^(*l*)^ is the nonlinear activation function at layer *l*, such as the ReLU, sigmoid, or tanh function.

ANNs are trained by minimizing a loss function–such as mean squared error in regression–via backpropagation and stochastic gradient descent (SGD) or adaptive optimization algorithms like Adam. During training, weights and biases are iteratively adjusted to reduce the prediction error. The flexibility of ANNs comes at the cost of increased computational demand and sensitivity to hyperparameter choices, such as learning rate, batch size, and architecture depth.

The universal approximation theorem guarantees that a sufficiently large neural network can approximate any continuous function on a compact domain to arbitrary accuracy. This makes ANNs powerful tools for modeling nonlinear ecological systems with high-dimensional interactions that may be difficult to capture with traditional parametric models.

To identify an optimal configuration for the neural network model, we performed a randomized hyperparameter search using 5-fold cross-validation. The search space included a range of architectural and training parameters, such as the number and size of hidden layers, dropout rates, learning rates, weight decay values, learning rate scheduling parameters (initial cycle length and minimum learning rate), and batch sizes. From a total of 648 possible configurations, five hyperparameter combinations were randomly sampled and evaluated. For each combination, five models were trained and validated across cross-validation folds, with mean squared error used as the evaluation metric. The average validation loss across 5-folds was used to determine the most effective configuration. This initial tuning phase provided a computationally efficient yet rigorous method for selecting a high-performing neural network architecture consistent with the evaluation procedures used for other models in the study.

The best-performing artificial neural network (ANN) configuration was selected based on average validation loss across five cross-validation folds. The selected hyperparameters follow.

- Hidden layer sizes (`hidden_sizes`): [**256, 128, 64**]
- Dropout rates per layer (`dropout_rates`): [**0.4, 0.3, 0.2**]
- Learning rate (`learning_rate`): **0.0005**
- Weight decay (`weight_decay`): **5e-5**
- Learning rate restart interval (`lr_restart_interval`): **20**
- Minimum learning rate (`min_lr`): **1e-6**
- Batch size (`batch_size`): **32**

## 5. Results

The comparative analysis revealed substantial performance differences among biological and ML models in predicting rainbow trout growth. Most models were similar in performance. Comparative analysis is provided after explication of all models.

### 5.1. Descriptive Statistics

Table 1 presents the key descriptive statistics for several numerical variables used in the analysis of Rainbow Trout growth and migration. The “Time at Large” variable, with a mean of 243.47 days and a standard deviation of 285.59, displays substantial variability, further confirmed by its high skewness (2.43) and kurtosis (6.92), indicating the presence of long-tailed observations and potential outliers. The physical growth measures, such as *L*_1_ and *L*_2_, show relatively symmetric distributions with low skewness and slightly negative kurtosis, suggesting a mild platykurtic shape. Environmental variables such as “Water Temperature” and “Solar Insolation” exhibit moderate variability and near-normal distributions, which may enhance model stability. Interestingly, “Release River Mile” is negatively skewed with an extremely high kurtosis (13.87), suggesting a heavily peaked distribution with frequent values near one end.

**Table 1:** Descriptive statistics of selected numerical variables.

| Variable | Mean | Std | Min | Median | Max | Skew | Kurtosis |
| --- | --- | --- | --- | --- | --- | --- | --- |
| Release River Mile | -3.74 | 0.69 | -14.89 | -3.75 | -1.96 | -1.12 | 13.87 |
| Time at Large | 243.47 | 285.59 | 28.00 | 130.50 | 2256.00 | 2.43 | 6.92 |
| Fork Length at Release ($L_1$) | 187.26 | 87.53 | 70.00 | 163.00 | 457.00 | 0.36 | -1.24 |
| Fork Length at Recapture ($L_2$) | 218.92 | 85.79 | 74.00 | 230.00 | 472.00 | 0.06 | -1.30 |
| Weight at Release | 118.78 | 132.94 | 3.50 | 51.00 | 1182.00 | 1.39 | 1.98 |
| Discharge | 18.65 | 4.14 | 11.33 | 17.81 | 32.11 | 1.34 | 1.74 |
| Water Temperature | 16.86 | 3.89 | 8.65 | 16.05 | 27.14 | 0.61 | 0.01 |
| Solar Insolation | 50.24 | 17.86 | 19.18 | 49.71 | 105.92 | 0.50 | 0.09 |
| Soluble Reactive $P$ Concentration | 0.01 | 0.00 | 0.00 | 0.01 | 0.02 | 0.42 | -0.27 |
| Rainbow Trout Biomass ( kg) | 12.38 | 6.09 | 5.56 | 10.45 | 34.03 | 0.99 | 0.20 |

### 5.2. Baseline Models

The baseline VBGM and Gompertz function were evaluated as mechanistic benchmarks, without the inclusion of covariates. The results indicate a clear performance difference between the two models. The baseline Gompertz model achieved a lower RMSE (61.52 mm) and mean absolute error (MAE = 49.81 mm) compared to the baseline VBGM (RMSE = 86.29 mm, MAE = 77.19 mm), suggesting superior predictive accuracy. Additionally, the Gompertz model explained a substantially higher proportion of variance in the test data, with an *R*^2^ of 0.492 and adjusted *R*^2^ of 0.483, whereas the VBGM performed no better than a mean-only model (*R*^2^ = 0.000). Information-theoretic criteria also favored the Gompertz formulation, which yielded substantially lower Akaike Information Criterion (AIC=32658.82), corrected AIC (32660.38), and Bayesian Information Criterion (BIC=32940.17) values than the VBGM (AIC = 34648.67, AICc = 34650.23, BIC = 34930.02). These results support the Gompertz model as the more effective baseline structure for capturing growth patterns in this dataset. Again, these models served as the comparative baseline.

### 5.3. Bayesian Models

The first three non-baseline model results were all Bayesian. In all cases, the convergence diagnostics indicated excellent mixing and stability across all parameters. In all cases, estimation was conducted using Markov Chain Monte Carlo (MCMC) with 4 chains and 1500 draws per chain (500 tuning, 1,000 posterior). The target accept=0.95 parameter increased the robustness of the NUTS in exploring complex posteriors.

The Gelman–Rubin statistic 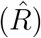 was 1.00 for all parameters across all models, demonstrating full convergence. Effective sample sizes (ESS) for both the bulk and tail of the posterior distributions were well above the commonly recommended threshold of 400, with many exceeding 6,000, indicating that the chains produced a large number of effectively independent samples. Moreover, no divergences or energy transition issues were reported for any of the models, further confirming their stability and reliability of the model.

#### 5.3.1. Fabens VBGM

The VBGM assumes that growth decelerates asymptotically as an organism approaches its theoretical maximum size *L*_∞_. The key strength of this model is that it provides posterior distributions over all unknowns, including individual-level growth rates, allowing for full uncertainty quantification. Unlike traditional VBGM approaches that assume constant growth rates, this model incorporates individual heterogeneity through environmental and temporal covariates, which is critical in ecological modeling where growth is often affected by dynamic habitat conditions.

The performance metrics for the VBGM suggest a solid, biologically informed baseline. The model yielded an RMSE of 16.82 and an MAE of 11.60, indicating that the average deviation of predicted fish lengths from observed values was around 11 to 17 mm. From a biological standpoint, this level of error is relatively modest given the natural variability in growth patterns among fish populations, and it suggests that the model is effectively capturing the underlying growth dynamics. The *R*^2^ value of 0.9620 and adjusted *R*^2^ of 0.9619 imply that over 96% of the variability in observed lengths is explained by the model, reinforcing the biological plausibility of VBGM in modeling somatic growth processes. Additionally, the AIC (24,945.04), AICc (24,945.05), and BIC (24,962.99) metrics indicate strong model parsimony, supporting the idea that the growth curve is both statistically and biologically well-specified without unnecessary complexity. (While a Widely-Applicable Information Criterion (WAIC) is generally preferred, use of the AIC, AICc, and BIC supported comparison.) Overall, these results affirm the VBGM’s capacity to capture length-at-age patterns with high fidelity, aligning with theoretical expectations of metabolic scaling and resource allocation in fish development.

#### 5.3.2. Gompertz Model

The Gompertz model also demonstrate reasonable but somewhat inferior predictive performance relative to the VBGM. The performance on the test set follows: RMSE=16.924, MAE=11.674, *R*^2^=0.962, Adjusted *R*^2^=0.962, AIC=24982.28, AICc=24982.29, BIC=25000.24.

While the Gompertz model outperformed the VGBM in the baseline, no-covariate setting likely due to its flexible curvature and early growth deceleration, its performance declined relative to VBGM once covariates were introduced in a Bayesian framework. One reason for this reversal is structural: the Gompertz model includes a single global rate parameter, which can limit its ability to accommodate feature-dependent variation in growth. In contrast, the VBGM, though initially more rigid, benefits more from the inclusion of biologically relevant covariates that can adjust parameters such as the asymptotic size or growth rate across individuals or conditions. This flexibility, combined with the regularizing effects of Bayesian estimation, enables the VBGM to better capture context-specific growth dynamics and reduces overfitting. The Gompertz model, by contrast, may become over-parameterized or less stable when extended with covariates, diminishing its initial advantage.

#### 5.3.3. Bayesian Linear Model

We fit a Bayesian linear regression model using *NumPyro* to estimate the relationship between trout recapture length (*L*_2_) and a combination of continuous and categorical predictors. All continuous predictors were standardized prior to model fitting, while the response variable (*L*_2_) remained in its original unit of millimeters to preserve ecological interpretability.

The model used a Normal prior for the coefficients (*β* ∼ 𝒩 (0, 5)), a Log-Normal prior on the standard deviation (log *σ* ∼ 𝒩 (0, 1)), and was sampled using the NUTS with 4 chains, each producing 1,500 posterior samples after a 500-sample tuning phase.

The Bayesian linear model demonstrated strong predictive accuracy compared to baseline methods. It achieved an *R*^2^ of 0.911, with an adjusted *R*^2^ of 0.910, indicating that the model captured the majority of the variance in the response variable. The RMSE was 25.72 mm, and the MAE was 18.54 mm, both reflecting substantial improvements over prior versions and competitive performance relative to more complex models.

The information criteria supported the model’s improved fit: AIC = 27,533.81, AICc = 27,535.44, and BIC = 27,821.15. These values are substantially lower than those observed in earlier specifications and within a competitive range compared to tree-based or boosting models, suggesting that the revised Bayesian linear model more effectively captured the underlying structure of the data.

While the Bayesian framework offers clear benefits in interpretability and uncertainty quantification, its application here did not result in a competitive model. Future improvements may involve incorporating hierarchical structures, interaction terms, or more flexible priors to better accommodate ecological complexity in trout growth modeling.

### 5.4. Trees Models

The results for each of the tree models follow. While traditional information criteria such as AIC, AICC, and BIC are typically derived from maximum likelihood estimation (MLE), many modern ML models (such as tree-based ensembles) do not possess closed-form likelihoods. In these cases, we approximate information criteria by assuming Gaussian residuals and computing the residual sum of squares (RSS) to estimate a pseudo-likelihood. Specifically, we treat the residuals as arising from a normal distribution with constant variance, and derive a log-likelihood under this assumption. The number of effective model parameters is approximated by counting the trainable parameters (e.g., via PyTorch’s `model.parameters()`), serving as a proxy for model complexity. While this approach does not provide true likelihood-based inference, it enables comparison across a diverse set of models using a consistent penalized error framework. As such, these criteria should be interpreted heuristically rather than as strict measures of statistical fit.

#### 5.4.1. RF

The tuned Random Forest model was then applied to the pristine test set data, achieving an RMSE of 18.40 mm and a MAE of 12.20 mm. The model explained approximately 95.45% of the variance in the outcome (*R*^2^ = 0.9545, adjusted *R*^2^ = 0.9538), indicating excellent predictive accuracy. The information criteria values (AIC = 25,563.22, AICc = 25,564.78, and BIC = 25,844.57) further confirm the model’s strong fit relative to other approaches. The three most important predictors identified by the Random Forest were *L*_1_ (length at release), time at large, and weight at release. These are intuitive and biologically plausible variables, reinforcing their relevance in predicting growth outcomes.

#### 5.4.2. XGBoost

XGBoost exhibited the lowest predictive error with an RMSE of 16.14 mm, slightly ahead of Random Forest (16.46 mm), VBGM (16.82 mm), and the Gompertz model (17.07 mm). Although the differences in RMSE are consistent across repeated validation folds, the absolute gap between the best-performing ML models and the Bayesian VBGM is under 1 mm. This suggests that the improvements in predictive accuracy, while statistically detectable, may not translate to biologically meaningful differences in growth estimation for management purposes.

#### 5.4.3. LightGBM

LightGBM achieved an RMSE of 16.25 mm and an MAE of 10.80 mm, delivering predictive performance nearly equivalent to XGBoost. With an *R*^2^ of 0.9645 and an adjusted *R*^2^ of 0.9640, the model explained a similarly high proportion of variance in the outcome. Although slightly higher than XGBoost, the information criteria (AIC = 24,830.76, AICc = 24,832.32, and BIC = 25,112.11) still reflect a strong model fit and reinforce LightGBM’s competitiveness among gradient boosting approaches. The top three most influential features were *L*_1_ (length at release), time at large, and the release river map, indicating the model’s ability to integrate both biological and spatial predictors effectively.

### 5.5. SVR Models

#### 5.5.1. Linear SVR

The linear SVR model showed noticeably weaker performance compared to the top ensemble methods in this study. It yielded an RMSE of 25.35 mm and a MAE of 15.40 mm–substantially higher than those of XGBoost (RMSE = 16.14) and LightGBM (RMSE = 16.25), and even lagging behind the Random Forest model (RMSE = 18.40). With an *R*^2^ of 0.9137 and adjusted *R*^2^ of 0.9122, the SVR explained just over 91% of the variance in recapture length–respectable, but clearly lower than the approximately 96% explained by the best-performing models.

From a model selection perspective, SVR also fared less favorably. Its AIC = 27,446.61 and BIC = 27,727.96 were substantially higher than those for XGBoost (AIC = 24,790.38, BIC = 25,071.73) and LightGBM (AIC = 24,830.76, BIC = 25,112.11), indicating poorer balance between model fit and complexity. It also trailed the Random Forest in information criteria, suggesting that SVR was less efficient overall in modeling the observed data.

While the linear Support Vector Regression (SVR) model did not achieve top-tier predictive performance compared to nonlinear models, it offered a distinct advantage in interpretability. Specifically, linear SVR provides direct coefficient estimates, enabling a clear understanding of the direction and relative magnitude of each input variableâĂŹs contribution to the predicted outcome. This directional insight is valuable for interpreting ecological or biological mechanisms in fish growth modeling. However, this transparency comes at the cost of flexibility, as the linear kernel assumes strictly additive and linear relationships between predictors and the response variable. Consequently, the model is unable to capture complex interactions or nonlinear effects, which likely explains its higher error metrics relative to kernel-based and ensemble models.

#### 5.5.2. Kernel-Optimized SVR

The RBF-kernel SVR model demonstrated strong predictive performance, outperforming several baseline and traditional models. It achieved a RMSE of 18.81, an MAE of 11.77, and an *R*^2^ value of 0.952. The adjusted *R*^2^ was similarly high at 0.951, indicating robust model fit even after accounting for model complexity. In terms of information-theoretic criteria, the RBF SVR produced an AIC of 25692.93, AICC of 25694.49, and BIC of 25974.28. While not the top-performing model across all metrics, it offered a balance between accuracy and generalization. Notably, it outperformed the linear SVR and Bayesian linear models across every metric, demonstrating the benefit of modeling nonlinear relationships in fish growth data through kernel-based methods.

To evaluate the relative influence of input features in the RBF SVR model, we employed permutation feature importance, a model-agnostic approach that quantifies importance based on changes in prediction error. This method involves systematically shuffling the values of each feature in the test set and measuring the resulting increase in the modelâĂŹs prediction error, thereby capturing how much each feature contributes to the modelâĂŹs performance. Results are incorporated to the feature importances analyses.

### 5.6. ANN

The Artificial Neural Network (ANN) model (see Figure 1) delivered solid predictive performance, with an RMSE of 18.47 mm and a MAE of 13.09 mm. These results placed it within the same general performance range as the Random Forest (RMSE = 18.40), though notably behind gradient-boosted models such as XGBoost (RMSE = 16.14) and LightGBM (RMSE = 16.25). The network achieved an *R*^2^ of 0.954 and an adjusted *R*^2^ of 0.952, indicating that it explained over 95% of the variance in recaptured length–strong, though slightly less than the top-performing boosting algorithms. While the ANN captured a meaningful portion of the underlying growth signal, its higher AIC (25,678.64), AICc (25,684.92), and BIC (26,241.34) suggest a trade-off in model efficiency compared to more parsimonious ensemble methods.

**Figure 1:**
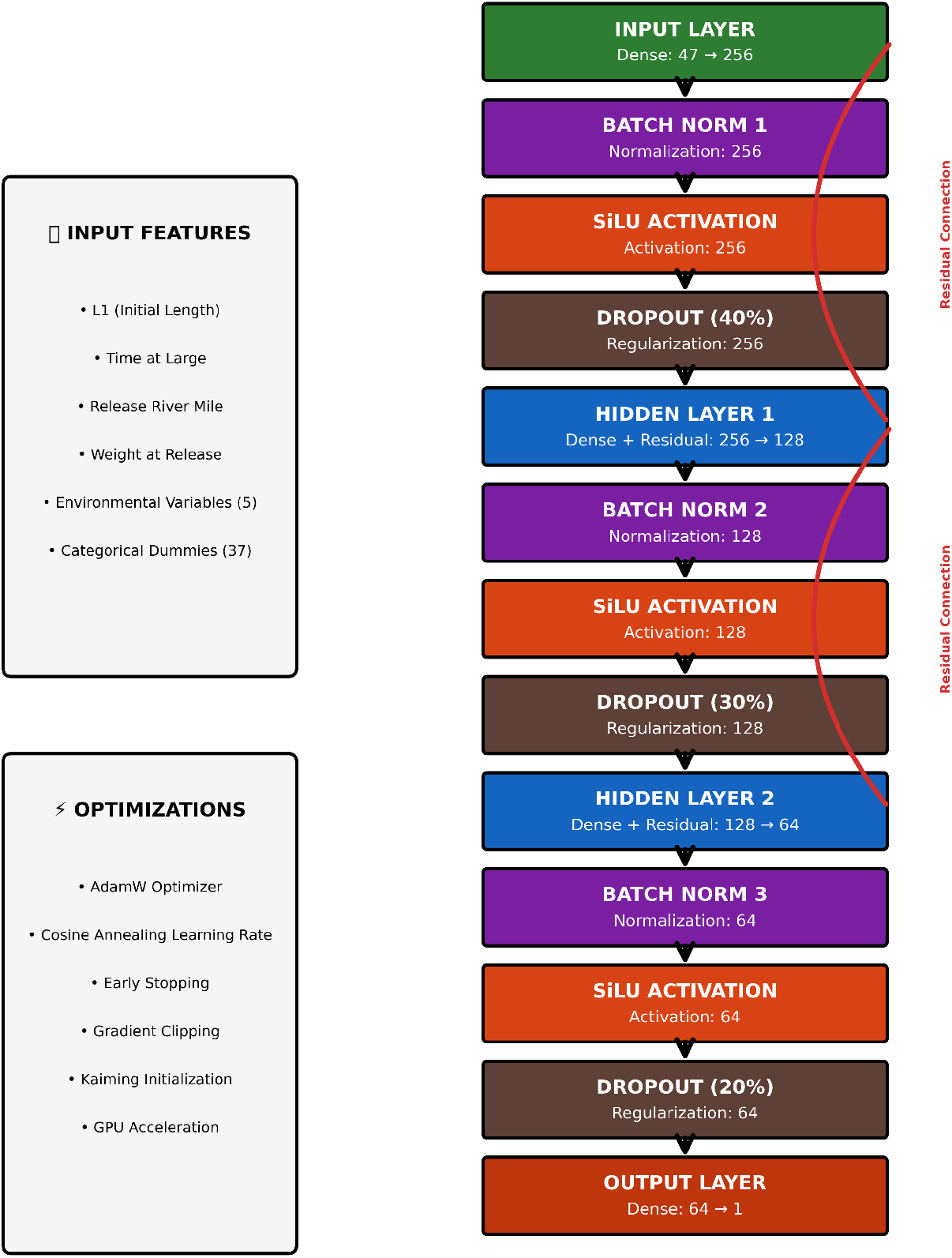
Neural Network Architecture

From an information-theoretic standpoint, the ANN yielded an AIC of 25,678.64 and a BIC of 26,241.34, both notably higher than those of the ensemble methods: XGBoost (AIC = 24,790.38, BIC = 25,071.73), Light-GBM (AIC = 24,830.76, BIC = 25,112.11), and Random Forest (AIC = 25,563.22, BIC = 25,844.57). These elevated values suggest a more pronounced penalty for model complexity, which is consistent with expectations given the parameter-rich architecture of neural networks. Unless effectively regularized, ANNs tend to be less parsimonious than tree-based methods. Integrated Gradients (IG), an attribution method that quantifies feature importance by integrating the gradients of the modelâĂŹs output with respect to its inputs along a straight-line path from a baseline (e.g., a zero vector) to the actual input, offer comparative insight into the internal prioritization of predictive factors by the network. IG are used for importance comparisons subsequently.

### 5.7. Ensembles

To synthesize the complementary strengths of biological and ML models, we implemented a stacked ensemble that combined predictions from the best-performing models within each paradigm as well as Bayesian Model Averaging (BMA). An explication follows.

For the first model, we used a linear regression model as a meta-learner to optimally weight predictions from the Bayesian VBGM and XGBoost. The meta-model was trained exclusively on out-of-sample predictions generated through cross-validation, ensuring that final ensemble weights were not influenced by data leakage. This stacking approach allowed the ensemble to adaptively balance interpretability and predictive flexibility, favoring one model or the other depending on local patterns in the data. By leveraging the structural insights of the biological model and the nonlinear predictive power of the ML model, the stacked ensemble achieved superior performance relative to all individual models. Its root mean square error (RMSE) was the lowest of any tested configuration, underscoring the practical value of integrating mechanistic understanding with data-driven learning in ecological forecasting.

The stacked ensemble, which combined predictions from the Bayesian VBGM and XGBoost, yielded the best overall performance across all evaluated models. It achieved the lowest RMSE (15.96 mm) and MAE (10.62 mm), indicating improved predictive accuracy over both its constituent models. The stacked ensemble also recorded the highest coefficient of determination (*R*^2^ = 0.9658) and adjusted *R*^2^ = 0.9658, reflecting its superior ability to explain variance in fork length growth while accounting for model complexity. From an information-theoretic perspective, it produced the lowest values for the Akaike Information Criterion (AIC = 24,635.91), corrected AIC (*AIC*_*C*_ = 24,635.92), and Bayesian Information Criterion (BIC = 24,653.87), further supporting its parsimony and goodness-of-fit relative to alternatives. These results confirm the effectiveness of the stacking strategy in integrating mechanistic and ML models, ultimately offering more accurate and robust growth forecasts than any single model alone.

The second approach, BMA, is a principled approach to model combination that accounts for uncertainty across competing models by averaging their predictions weighted by their posterior probabilities. Unlike traditional ensemble methods that may use equal or heuristic weights, BMA incorporates Bayesian reasoning to assign greater influence to models that explain the data well while penalizing overly complex or poorly fitting models. This results in predictive distributions that naturally reflect model uncertainty and often lead to improved generalization performance.

The results for the Bayesian Model Averaging (BMA) ensemble reveal its strength as a competitive and balanced forecasting approach. In terms of RMSE and MAE, BMA matches the performance of XGBoost exactly— achieving an RMSE of 16.137 and an MAE of 10.722—indicating that its averaged predictions closely align with those of the strongest individual model. However, BMA information criteria values are uniformly lower than or comparable to other models, notably outperforming stacked ensembles, which although slightly better in RMSE, incur higher complexity penalties. This reflects BMAâĂŹs Bayesian advantage: it combines predictive strength with parsimony, minimizing overfitting risk by integrating model uncertainty. Overall, BMA delivers a robust, interpretable alternative to black-box models by leveraging posterior-weighted synthesis while maintaining statistical efficiency.

### 5.8. Model Comparison

Model comparisons across metrics and coefficients identify both diverging and converging models. Explication follows.

#### 5.8.1. Metrics

Table 2 summarizes the performance of all models using RMSE, MAE, *R*^2^, adjusted *R*^2^, and information criteria (AIC, AICC, BIC). The best overall performance was achieved by the *Stacked Ensemble*, which outperformed all individual models with the lowest RMSE (15.96 mm), lowest MAE (10.62 mm), and highest *R*^2^ (0.9658). This highlights the effectiveness of aggregating predictive strengths from both mechanistic and ML paradigms.

**Table 2:**
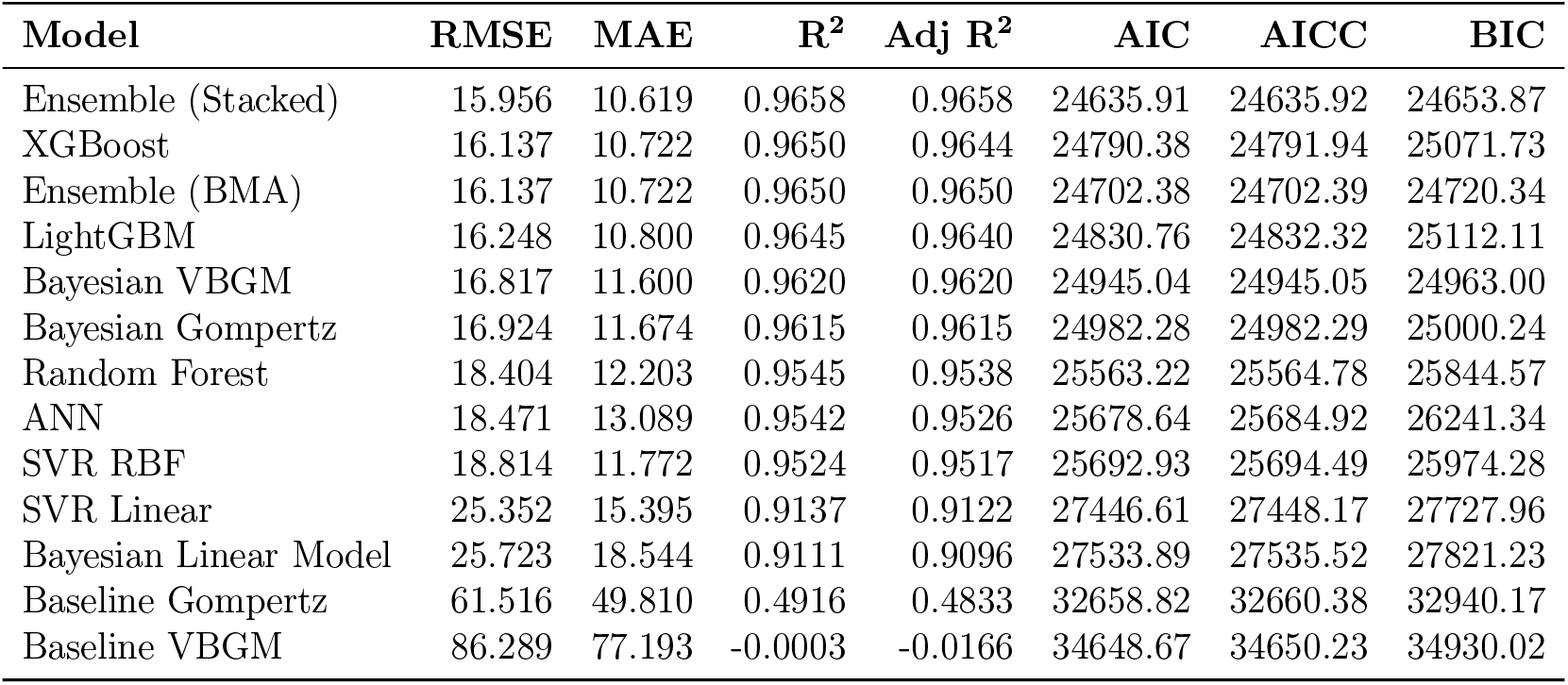
Comparison of Model Performance Metrics (Sorted by RMSE)

| Model | RMSE | MAE | R<sup>2</sup> | Adj R<sup>2</sup> | AIC | AICC | BIC |
| --- | --- | --- | --- | --- | --- | --- | --- |
| Ensemble (Stacked) | 15.956 | 10.619 | 0.9658 | 0.9658 | 24635.91 | 24635.92 | 24653.87 |
| XGBoost | 16.137 | 10.722 | 0.9650 | 0.9644 | 24790.38 | 24791.94 | 25071.73 |
| Ensemble (BMA) | 16.137 | 10.722 | 0.9650 | 0.9650 | 24702.38 | 24702.39 | 24720.34 |
| LightGBM | 16.248 | 10.800 | 0.9645 | 0.9640 | 24830.76 | 24832.32 | 25112.11 |
| Bayesian VBGM | 16.817 | 11.600 | 0.9620 | 0.9620 | 24945.04 | 24945.05 | 24963.00 |
| Bayesian Gompertz | 16.924 | 11.674 | 0.9615 | 0.9615 | 24982.28 | 24982.29 | 25000.24 |
| Random Forest | 18.404 | 12.203 | 0.9545 | 0.9538 | 25563.22 | 25564.78 | 25844.57 |
| ANN | 18.471 | 13.089 | 0.9542 | 0.9526 | 25678.64 | 25684.92 | 26241.34 |
| SVR RBF | 18.814 | 11.772 | 0.9524 | 0.9517 | 25692.93 | 25694.49 | 25974.28 |
| SVR Linear | 25.352 | 15.395 | 0.9137 | 0.9122 | 27446.61 | 27448.17 | 27727.96 |
| Bayesian Linear Model | 25.723 | 18.544 | 0.9111 | 0.9096 | 27533.89 | 27535.52 | 27821.23 |
| Baseline Gompertz | 61.516 | 49.810 | 0.4916 | 0.4833 | 32658.82 | 32660.38 | 32940.17 |
| Baseline VBGM | 86.289 | 77.193 | -0.0003 | -0.0166 | 34648.67 | 34650.23 | 34930.02 |

Among individual learners, *XGBoost* and *LightGBM* performed exceptionally well, achieving RMSE values of 16.14 mm and 16.25 mm, respectively, with high *R*^2^ values and favorable information criteria. These results reaffirm the ability of gradient boosting algorithms to model complex non-linear processes in ecological systems. Bayesian biological models, including the *VBGM* and the *Gompertz equation*, also ranked competitively, demonstrating that biologically grounded models remain viable, especially when well-calibrated, informed by data, and inclusive of covariates.

In contrast, the *Baseline VBGM* and *Baseline Gompertz* modelsâĂŤfitted without covariatesâĂŤperformed poorly, with RMSE values of 86.29 mm and 61.52 mm, respectively, and substantially lower *R*^2^ values. These results emphasize the limitations of using biologically motivated models without accounting for relevant environmental and contextual variables, reinforcing the importance of covariate inclusion for predictive accuracy.

Outside of the poor-performing baseline models, the *Bayesian Linear Model* and *Linear SVR* showed the weakest performance among fully trained models, with RMSEs exceeding 25 mm and comparatively lower *R*^2^ values, indicating a poor fit to the data. However, the *Radial Basis Function (RBF) SVR* improved substantially upon the linear variant, achieving an RMSE of 18.81 mm and *R*^2^ = 0.952, illustrating the benefit of nonlinear kernel methods in capturing more complex relationships. The *Random Forest* and *Neural Network* models also produced *R*^2^ values above 0.95, but incurred higher information criteria, reflecting greater model complexity and possibly reduced generalization.

These findings support a natural next step: applying *Bayesian Model Averaging (BMA)* to integrate top-performing models across paradigms. The BMA approach achieved similar accuracy to XGBoost while explicitly incorporating model uncertainty through posterior-weighted predictions. This technique may be particularly advantageous in ecological forecasting contexts where interpretability, uncertainty quantification, and generalizability are critical. Future research could assess whether BMA further enhances model robustness while preserving the domain-specific strengths of both mechanistic and ML-based approaches.

#### 5.8.2. Coefficients

Figure 2 illustrates the top nine continuous features identified through average normalized importance scores across all models. The feature importance scores were normalized using min-max scaling to a [0,1] range to enable direct comparison across different models and metrics. The variable *L*_1_, representing asymptotic length, emerged as the most influential predictor overall. Its importance was consistently high across nearly all modeling approaches, with particularly strong influence in the Bayesian Gompertz, Bayesian VBGM, and SVR RBF models. This is expected, as *L*_1_ is a core parameter in both the VBGM and the Gompertz models, where it serves as a theoretical maximum size. These mechanistic models explicitly rely on this variable to define growth trajectories, which explains its critical importance.

**Figure 2:**
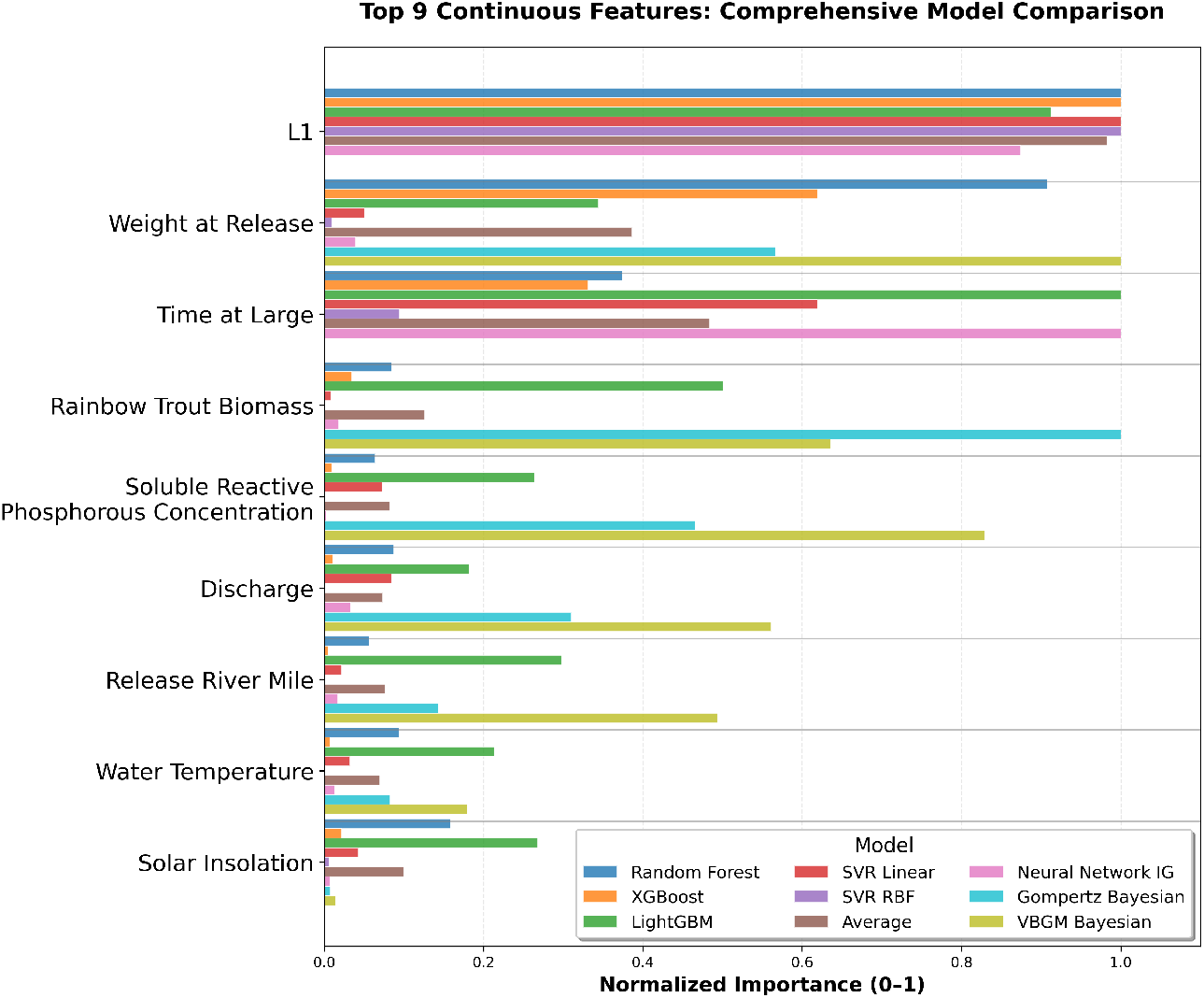
Normalized Importance Metrics

The variable *Time at Large*, another essential parameter in traditional growth models, also ranked highly in terms of predictive contribution. In both the VBGM and Gompertz frameworks, *Time at Large* maps directly to the growth curve’s temporal component, linking observed size back to age or exposure duration. Its strong influence in ML models (including LightGBM and Neural Networks) underscores its general predictive relevance, even outside mechanistic contexts.

Additional high-ranking features included *Weight at Release, Rainbow Trout Biomass*, and *Soluble Reactive Phosphorous Concentration*. These environmental or biological covariates were less central in traditional growth models but proved informative in ensemble methods like XGBoost and Random Forest. This suggests that data-driven models may capture interaction effects and context-dependent relationships that lie outside the scope of classical equations.

The consistency of *L*_1_ and *Time at Large* across both mechanistic and ML approaches validates their foundational role in growth modeling. Meanwhile, the variable-specific strengths of different algorithms highlight the complementary nature of ensemble and domain-specific methods when predicting ecological phenomena.

### 5.9. Bayesian Probabilistic Comparison of RMSE

Pairwise posterior probabilities were computed from the joint posterior distribution over model RMSEs to estimate the probability that one model outperformed another, as summarized in Table 3. These values represent the proportion of posterior samples for which the RMSE of one model was lower than that of another.

**Table 3:** Pairwise Stochastic Dominance Probabilities (Sorted by Number of Models Beaten)

| Model | Ensemble (Stacked) | LightGBM | XGBoost | Ensemble (BMA) | Bayesian VBGM | Bayesian Gompertz | Random Forest | ANN | SVR_Radial | SVR_Linear | Bayesian Linear |
| --- | --- | --- | --- | --- | --- | --- | --- | --- | --- | --- | --- |
| Ensemble (Stacked) | - | 0.512 | 0.509 | 0.506 | 0.525 | 0.535 | 0.589 | 0.596 | 0.612 | 0.771 | 0.776 |
| LightGBM | 0.488 | - | 0.501 | 0.501 | 0.524 | 0.529 | 0.585 | 0.583 | 0.598 | 0.770 | 0.775 |
| XGBoost | 0.491 | 0.499 | - | 0.504 | 0.527 | 0.543 | 0.583 | 0.593 | 0.598 | 0.765 | 0.775 |
| Ensemble (BMA) | 0.494 | 0.498 | 0.496 | - | 0.528 | 0.530 | 0.582 | 0.590 | 0.607 | 0.766 | 0.774 |
| Bayesian VBGM | 0.475 | 0.476 | 0.473 | 0.471 | - | 0.508 | 0.562 | 0.568 | 0.577 | 0.755 | 0.759 |
| Bayesian Gompertz | 0.465 | 0.471 | 0.457 | 0.470 | 0.491 | - | 0.550 | 0.569 | 0.580 | 0.756 | 0.758 |
| Random Forest | 0.411 | 0.415 | 0.417 | 0.418 | 0.438 | 0.450 | - | 0.503 | 0.520 | 0.724 | 0.727 |
| ANN | 0.404 | 0.417 | 0.407 | 0.410 | 0.432 | 0.431 | 0.497 | - | 0.519 | 0.722 | 0.724 |
| SVR_Radial | 0.388 | 0.402 | 0.402 | 0.393 | 0.423 | 0.420 | 0.480 | 0.480 | - | 0.715 | 0.726 |
| SVR_Linear | 0.229 | 0.230 | 0.235 | 0.234 | 0.245 | 0.244 | 0.276 | 0.279 | 0.285 | - | 0.514 |
| Bayesian Linear | 0.224 | 0.225 | 0.225 | 0.226 | 0.241 | 0.241 | 0.273 | 0.276 | 0.274 | 0.486 | - |

## 6. Discussion

This study aimed to bridge traditional ecological modeling and advanced ML (ML) methods to forecast invasive rainbow trout growth in the Lower Colorado River, highlighting the complementary strengths and trade-offs inherent in each modeling paradigm. Our findings are contextualized within existing literature, emphasizing both ecological theory and methodological advancements.

The most striking finding is the transformative impact of incorporating covariates and advanced modeling approaches. Compared to baseline models without covariates, our best-performing methods achieved dramatic improvements: the stacked ensemble reduced RMSE by 70.33 mm (81.5% reduction) relative to the baseline VBGM and by 45.56 mm (74.1% reduction) relative to the baseline Gompertz model. These substantial gains underscore the critical importance of environmental and biometric covariates in biological growth modeling and demonstrate the severe limitations of traditional approaches that ignore ecological context.

The predictive results revealed important insights into model effectiveness for this ecological application. The stacked ensemble achieved the best point estimate performance (RMSE = 15.96 mm, *R*^2^ = 0.9658) and demonstrated stochastic dominance by outperforming all other models in the majority of posterior samples. This result supports the effectiveness of hybrid approaches that integrate mechanistic structure with data-driven flexibility (Li et al., 2021), illustrating that ensemble strategies can deliver both high average accuracy and consistent superiority across the full distribution of outcomes.

LightGBM (RMSE = 16.25 mm) and XGBoost (RMSE = 16.14 mm) formed a competitive second tier, with LightGBM achieving 9 dominances and XGBoost achieving 8 dominances in the stochastic analysis. Both models achieved over 80% RMSE reductions compared to baseline approaches, reinforcing the particular suitability of gradient boosting methods for ecological modeling, as demonstrated by Ho and Goethals (2022), who showcased ML flexibility in capturing complex, nonlinear environmental interactions in freshwater ecosystems. Our findings extend this work by showing that tree-based ensemble methods offer robust performance characteristics for biological growth prediction, though the stacked ensemble approach provides additional advantages through model integration.

The Bayesian Model Averaging (BMA) ensemble achieved competitive performance (RMSE = 16.14 mm, *R*^2^ = 0.9650) while demonstrating 7 dominances in the stochastic analysis. BMA’s capacity to average across models weighted by their posterior evidence presents a compelling alternative that balances predictive performance with explicit uncertainty quantification, especially valuable in ecological contexts where robust inference and model uncertainty assessment are paramount (Flores et al., 2024). The BMA approach also achieved an 81.3% RMSE reduction compared to the baseline VBGM, demonstrating that principled model averaging can achieve transformative performance improvements while maintaining probabilistic rigor.

Importantly, our analysis reaffirms the continued relevance of traditional growth models such as the VBGM when enhanced with covariates and Bayesian estimation, though with important caveats about their predictive consistency relative to modern approaches. The Bayesian VBGM achieved competitive point estimate performance (RMSE = 16.82 mm) and maintains crucial biological interpretability through meaningful parameters including asymptotic size and intrinsic growth rates (von Bertalanffy, 1938; Fabens, 1965). The model achieved 6 dominances in the stochastic analysis, demonstrating solid middle-tier performance while delivering a 80.5% RMSE improvement over its baseline counterpart. However, it was consistently outperformed by ensemble methods and leading ML algorithms, suggesting that while mechanistic models provide essential ecological insights for fisheries management and theoretical understanding, their predictive reliability may be more variable than that of advanced data-driven approaches.

The comparative analysis highlighted critical differences between mechanistic and ML models regarding both predictive consistency and interpretability. The top-performing ensemble and ML methods demonstrated exceptional ability to capture subtle environmental and biometric interactions while maintaining robust performance across diverse conditions. However, their reliance on feature importance metrics, rather than explicit biological parameters, limits direct ecological inference. This aligns with critiques in the literature emphasizing the interpretive challenges associated with ML approaches (Pineda-Metz et al., 2023; Bhai et al., 2024). Conversely, mechanistic models such as VBGM offer clearer connections to biological theory, supporting the ecological interpretations required for informed management decisions (Fabens, 1965), even if their predictive distributions show greater variability relative to leading ensemble methods.

The success of the stacked ensemble approach demonstrates the potential for hybrid modeling frameworks that integrate the complementary strengths of different paradigms. By combining predictions from XGBoost and the VBGM, the stacked ensemble achieved both the best point estimate performance and the highest dominance count, suggesting that such integration can harness the predictive power of ML while incorporating the biological grounding of mechanistic models. This represents a promising direction for ecological forecasting where both accuracy and interpretability are valued.

The Bayesian probabilistic comparison provided a more comprehensive performance assessment than traditional point estimate metrics alone. Using posterior distributions over model RMSEs, we computed pairwise dominance probabilities that revealed important performance patterns not apparent from RMSE values alone. The stacked ensemble’s complete dominance across all pairwise comparisons strengthens confidence in its selection for practical applications, while the competitive showing of individual ML methods and BMA demonstrates the value of multiple modeling approaches for different ecological contexts.

Support Vector Regression models showed variable performance in both point estimates and probabilistic evaluation. SVR Radial achieved moderate point estimate performance (RMSE = 18.81 mm) but managed only 2 dominances in the stochastic analysis, while SVR Linear performed poorly across both metrics. However, even the weaker-performing SVR models achieved substantial improvements over baseline approaches (78.2% and 70.6% RMSE reductions, respectively), suggesting that kernel-based methods, while less optimal than tree-based ensemble approaches, still provide meaningful advantages when covariates are properly incorporated.

The systematic evaluation revealed that predictive performance in ecological modeling benefits from comprehensive assessment frameworks that consider both point estimates and distributional characteristics. The alignment between RMSE rankings and stochastic dominance patterns for the top-performing models provides confidence in the robustness of these findings, while the ensemble methods’ superior performance across both evaluation frameworks strengthens their candidacy for practical applications.

Furthermore, the identification of critical predictors such as initial length (*L*_1_), time at large, and weight at release through ML methods aligns with well-established ecological theories of fish growth regulation (Sumpter, 1991). Our findings reinforce the central role of intrinsic and environmental factors previously documented in rainbow trout literature, confirming the robustness and ecological plausibility of both individual ML algorithms and ensemble approaches in capturing these fundamental biological relationships.

The limitations in our dataset, notably the absence of consistent individual identifiers across capture events, required treating each capture event independently. This constraint, while mitigated by our modeling strategies, underscores the need for improved tagging protocols and longitudinal study designs, as emphasized by Korman and Yard (2017). The lack of individual tracking limited our ability to apply repeated-measures or hierarchical modeling frameworks that could more fully leverage temporal dependencies and individual growth trajectories. Additionally, while environmental covariates were included, they were derived from reach-level estimates and did not always reflect the microhabitat conditions experienced by individual fish. This spatial aggregation may have introduced noise into model inputs, particularly for variables like temperature and discharge that vary significantly within short river segments. Our modeling framework was also developed within the unique hydrological and ecological context of the Lower Colorado River, and its transferability to other systems with different flow regimes, food webs, or stocking practices remains an open question. Lastly, while the sample size was robust overall, growth observations were disproportionately concentrated in certain size classes and seasons, potentially introducing bias in model learning and performance across the full life cycle. Future studies incorporating higher-resolution environmental data, balanced seasonal sampling, and repeated-capture designs with individual tracking would provide a stronger foundation for both mechanistic and ML models, further improving accuracy, uncertainty quantification, and biological interpretability.

Overall, this study supports a nuanced view wherein ensemble methods that combine mechanistic and ML models offer superior performance characteristics compared to individual approaches. While ML methods excel in predictive accuracy and flexibility, and traditional growth models remain essential for ecological interpretation and theoretical grounding, the integration of these approaches through ensemble frameworks provides the most robust solution for ecological forecasting applications. The success of both stacked ensembles and BMA demonstrates multiple pathways for achieving this integration, with the choice depending on specific application requirements for prediction accuracy versus uncertainty quantification.

The performance improvements demonstrated by covariate-enhanced modeling approaches are substantial and biologically meaningful. Achieving 70-80% RMSE reductions compared to baseline models represents a transformative advance in growth prediction capability, with improvements of 45-70 mm representing 20% to 32% of mean fish length in this study. These gains significantly exceed typical measurement precision and natural growth variability, providing ecologically and management-relevant improvements in forecasting reliability. The additional modest improvements (0.2-0.9 mm RMSE reduction) achieved by ensemble methods over individual covariate-based models, while smaller in absolute terms, come with significantly enhanced consistency across diverse conditions, offering further value for applications where predictive reliability is critical.

## 7. Conclusion

This study demonstrates the exceptional value of ensemble methods that integrate traditional biological models with ML techniques for ecological forecasting, while revealing the transformative impact of incorporating environmental and biometric covariates in growth modeling. Among all models tested, the stacked ensemble emerged as the top performer, achieving both the best point estimate performance (RMSE = 15.96 mm, *R*^2^ = 0.9658) and stochastic dominance over all other covariate-based models. This hybrid framework, combining predictions from XGBoost and the von Bertalanffy growth model (VBGM), illustrates how the predictive power of advanced ML can be grounded in biological realism.

The most significant finding is the substantial improvement achieved through covariate inclusion and advanced modeling techniques. Top-performing models showed 70 to 80 percent RMSE reductions relative to baseline approaches without covariates, with gains of 45 to 70 mmâĂŤequivalent to 20 to 32 percent of mean fish length. These improvements are not incremental; they reflect a fundamental shift in forecasting capability enabled by incorporating environmental context and model complexity.

The modeling comparison revealed a clear performance hierarchy. Gradient boosting methods like LightGBM (9 dominances) and XGBoost (8 dominances) formed a strong second tier, providing highly accurate and consistent performance. Bayesian Model Averaging (7 dominances) offered competitive accuracy and unique value through probabilistic weighting and uncertainty quantification. Mechanistic models such as the Bayesian VBGM, while not top-ranked, still delivered substantial performance improvements and remain indispensable for ecological interpretation, especially regarding growth ceilings, maturation thresholds, and response to flow regimes.

The combined use of point-estimate metrics and stochastic dominance analysis strengthened confidence in model robustness. Stochastic dominance revealed that ensemble approaches not only reduce error but outperform alternative models across the entire posterior distribution, which is critical for applications under uncertainty. These results support comprehensive evaluation methods that move beyond single-metric comparisons and emphasize distributional consistency.

From a management perspective, these findings carry important operational implications. First, managers should invest in collecting and integrating environmental covariatesâĂŤparticularly water temperature, flow regime, and photoperiodâĂŤalongside biometric data. These variables consistently improved model performance and offer actionable levers for management intervention. Second, stacked ensemble models should be used for forecasting population dynamics when accurate prediction is critical to decision-making, such as evaluating stocking schedules or predicting growth under altered dam release regimes. Third, mechanistic models should be maintained within ensemble frameworks to preserve interpretability and support biologically informed policy. Their parameters can guide flow thresholds, fishing quotas, and habitat restoration targets even if not optimal alone for prediction. Finally, for regulatory scenarios requiring risk estimates or management under uncertainty, Bayesian Model Averaging provides an effective tool for quantifying predictive distributions and informing precautionary decisions.

The integrative framework demonstrated hereâĂŤunifying mechanistic understanding, modern ML, and probabilistic evaluationâĂŤoffers a generalizable methodology for ecological forecasting. It is well-suited for complex biological systems where both accuracy and interpretability are essential. Future research should refine methods of ensemble weighting, assess model generalizability across species and systems, and further explore policy-aligned metrics that link prediction to management thresholds. Ecological managers are encouraged to adopt these hybrid strategies not as theoretical enhancements, but as practical tools capable of supporting evidence-based conservation in dynamic and uncertain environments.

## 8. Glossary

Artificial Neural Network (ANN): A computational model inspired by the brain, capable of learning non-linear data relationships
Bayesian Inference: A statistical approach that updates prior beliefs based on observed data
Cross-Validation (CV): A method to assess model performance using repeated data partitioning
Δ*t*: Time elapsed between initial and final measurements
Gradient Boosting (GB): A technique that builds models sequentially to minimize the residual errors of previous models
Hyperparameter Tuning: The process of optimizing algorithm settings to enhance performance
*K* (Growth Coefficient): Rate parameter in the VBGM indicating how quickly the organism approaches its maximum size
*L*_1_: Measured length of a fish at its first observation or tagging
*L*_2_: Predicted length of a fish at a later time point
*L*_∞_: Theoretical maximum length an organism can reach
Machine Learning (ML): Algorithms that learn patterns from data to make predictions without explicit instructions
Mean Absolute Error (MAE): The average absolute difference between predicted and observed values
No-U-Turn Sampler (NUTS): An adaptive Hamiltonian Monte Carlo algorithm that automatically tunes step size and path length to efficiently sample from complex posterior distributions without manual parameter tuning
Overfitting: A modeling error where the model captures noise instead of the underlying pattern
Predictive Congruence: The degree to which different models produce similar or equivalent predictions
Predictive Modeling: The use of statistical or machine learning methods to forecast outcomes from data
Random Forest (RF): An ensemble learning method using multiple decision trees to improve prediction accuracy
Root Mean Square Error (RMSE): A standard measure of prediction error magnitude in models
Support Vector Regression (SVR): A machine learning method for predicting continuous outcomes using margin-based optimization
Tag-Recapture Study: A method to estimate growth or survival by tagging individuals and recapturing them later
von Bertalanffy Growth Model (VBGM): A biological growth function describing the length of an organism as a function of age, commonly used in fisheries science

## 9. Declarations

### 9.1. Use of Generative Artificial Intelligence

During the preparation of this work the authorsused ChatGPT4o for editing content. After using this tool/service, the authors reviewed and edited the content as needed and take full responsibility for the content of the published article.

During the preparation of this work the authors used Claude.ai for code debugging. After using this tool/service, the authors reviewed and edited the content as needed and take full responsibility for the content of the published article.

### 9.3. Data Availability

The underlying USGS datasets are publicly available as cited. Analysis code and processed datasets will be made available in a public repository upon publication to ensure full reproducibility.

## 10. Author Contributions

P.L.: Conceptualization, Methodology, Software, Writing Original Draft. L.F.: Methodology, Software, Supervision, Writing Original Draft.

